# A two-process model of *Drosophila* sleep reveals an inter-dependence between circadian clock speed and the rate of sleep pressure decay

**DOI:** 10.1101/2022.08.12.503775

**Authors:** Lakshman Abhilash, Orie Thomas Shafer

## Abstract

Sleep is controlled by two processes – a circadian clock that regulates its timing and a homeostat that regulates the drive to sleep. *Drosophila* has been an insightful model for understanding both processes. For four decades, Borbély and Daan’s two-process model has provided a powerful framework for understanding how circadian and homeostatic processes regulate sleep. However, the field of fly sleep has not employed such a model as a framework for the investigation of sleep. To this end, we have adapted the two-process model to the fly and establish its utility by showing that it can provide empirically testable predictions regarding the circadian and homeostatic control of fly sleep. We show that the ultradian rhythms previously reported for loss-of-function clock mutants are a predictable consequence of a functional sleep homeostat in the absence of a functioning circadian system. We find that a model in which the circadian clock speed and homeostatic rates act without influencing each other provides imprecise predictions regarding how clock speed influences the strength of sleep rhythms and the amount of daily sleep. We also find that quantitatively good fits between empirical values and model predictions were achieved only when clock speeds were positively correlated with rates of decay of sleep pressure. Our results indicate that longer sleep bouts better reflect the homeostatic process than the current definition of sleep as any inactivity lasting five minutes or more. This two-process model represents a powerful framework for future work on the molecular and physiological regulation of fly sleep.

## Introduction

Sleep-like states are ubiquitous within the animal kingdom [1] and the complexity of sleep behavior has prompted many attempts to mathematically model its regulation. Such models provide a powerful quantitative framework for the experimental investigation of sleep regulation across taxa. They are particularly useful by virtue of their provision of testable hypotheses [2–9]. Among the most successful models of sleep regulation are variants of Borbély’s two-process model [2,4–7], which posits the presence of a sleep pressure that builds during wakefulness (Fig. 1A). Once this sleep pressure reaches an upper threshold, sleep is induced. During sleep, pressure falls until it reaches a lower threshold, at which point wakefulness is initiated. According to this model, the build-up and discharge of sleep pressure is driven by one process – the sleep homeostatic component or the S-process, whereas the upper and lower thresholds change predictably throughout the day and are set by a second process– the circadian clock or the C-process (Fig. 1A). Though myriad factors impact sleep [10], the regulation of typical sleep/wake cycles can be explained by just these two processes acting in concert but operating separately, that is, the S-process can operate without a functional C-process and *vice versa* [6,7].

**Figure 1:**
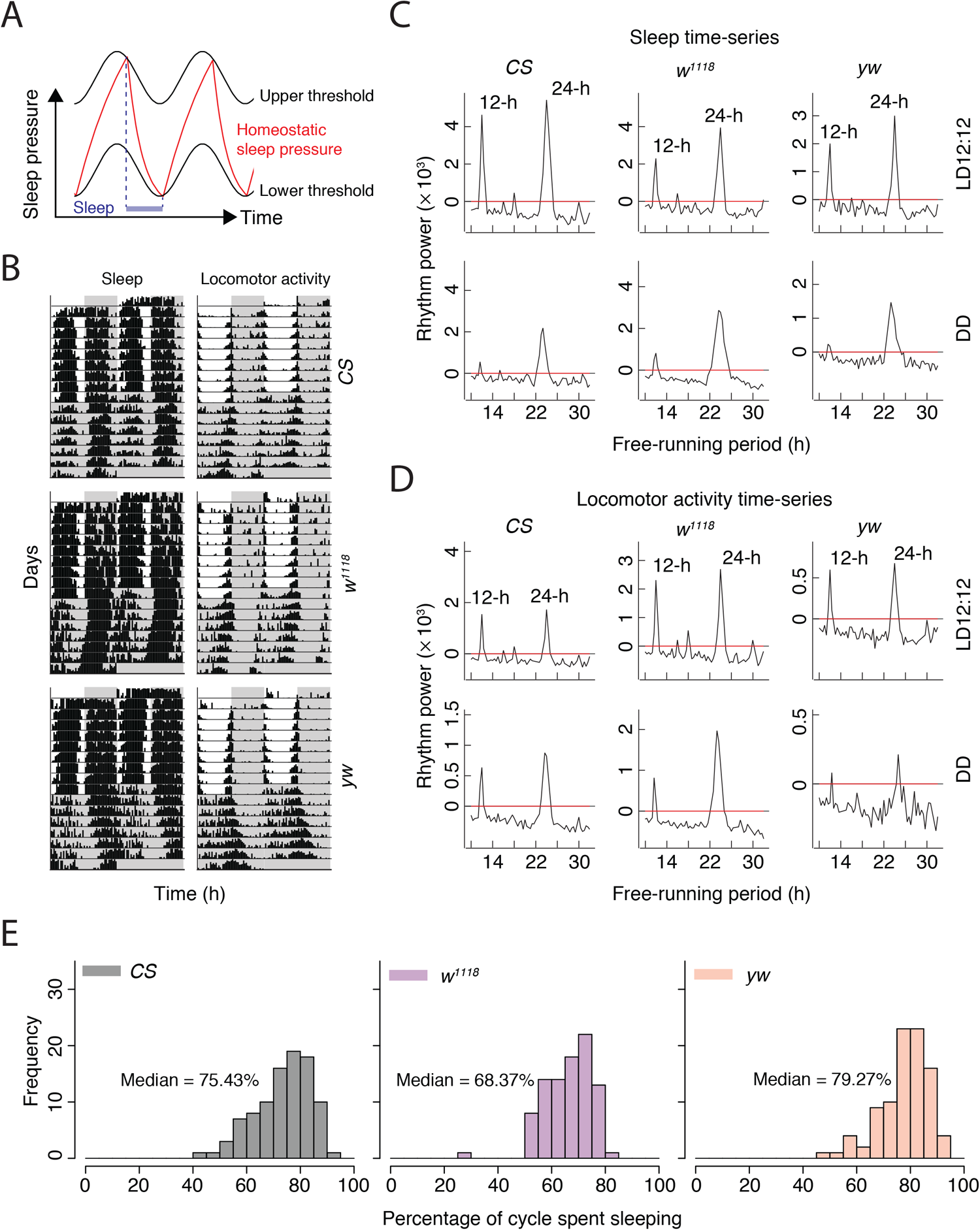
Circadian analysis of *Drosophila* sleep. (A) A schematic of a two-process model that leads to one episode of sleep/cycle. (B) Somnograms (left) and actograms (right) of *Canton-S* (CS; top), *w^1118^* (middle), and *yw* (bottom) flies under light/dark 12:12 (LD12:12) cycles and constant darkness (DD). Gray shaded regions indicate the dark phase of the LD cycle. Representative Chi-square periodograms of sleep time-series (C) under LD12:12 (top) and DD (bottom), and of locomotor activity time-series (D) under LD12:12 (top) and DD (bottom) of the three genotypes of flies. The red lines indicate the Chi-square significance line at *α* = 0.05. Height of peaks of the black lines indicate the strength of that periodicity in the time-series. (E) Frequency distributions of percentage time spent sleeping/24-h cycle in the three genotypes under LD12:12.

The two-process model has provided a powerful conceptual framework in the field for nearly four decades owing to its ability to explain changes in slow wave sleep and predict the timing and amount of sleep, particularly in humans, both during normal sleep/wake cycles and in the aftermath of sleep deprivation [5–7]. The model has also been used over the years to inform sleep clinicians in the treatment of sleep disorders associated with major depression, seasonal affective disorders, irregular sleep patterns, and the application of bright light therapy [7]. Additionally, the model provides a powerful, quantitative framework for assessing interactions between the two processes and how they might contribute to the control of sleep [11]. Despite the longstanding utility of the two-process model for understanding human sleep, it has been underutilized as a quantitative platform for understanding the regulation of sleep in model systems amenable to the genetic and molecular manipulation of the brain.

Though *Drosophila* has been a critical model system for understanding the genetic, neuronal and physiological bases of circadian rhythms since the 1970s, it has become an informative model for understanding sleep only over the past two decades [12–14]. Furthermore, because of the availability of a powerful array of genetic tools and the ability to regulate gene expression spatially and temporally in the fly, the *Drosophila* sleep field has made significant advances in understanding the regulation of sleep [15–17]. While much of the focus of fly sleep research has been on the identification of genes and neural networks that regulate the amount of sleep [14,16,18–27] relatively little is known about the regulation of sleep timing (e.g., [28]). Therefore, much remains to be learned about how circadian clocks and homeostatic control mechanisms interact to regulate multiple aspects of sleep.

While there are significant gaps in our understanding of sleep regulation, the fly continues to provide insights into its mechanisms [14]. A major reason for this is the apparent similarity (behavioral, genetic, and physiological) between fly and mammalian sleep [29]. For example, as in mammals, flies display bouts of extended inactivity characterized by increased arousal thresholds, rapid reversibility, and homeostatic rebound in response to sleep deprivation [12,15,30]. In addition, a major feature of mammalian sleep is the occurrence of distinct stages of sleep, i.e., rapid eye movement (REM) and several non-REM (NREM) sleep stages [31]. Although various durations of inactivity have previously been classified as distinct epochs of sleep [12,32], fly sleep is now almost universally treated as a unitary state, with any period of inactivity lasting more than five minutes being considered a sleep like state [14,32]. However, recent work has provided strong evidence for multiple sleep stages in flies, with longer bouts of inactivity representing physiologically, metabolically, and behaviorally distinct sleep states compared to shorter bouts [33–39], thereby adding an additional degree of similarity in sleep between flies and mammals.

Owing to the predictive power of the two-process model of human sleep regulation, we sought to establish a quantitative theoretical framework for a two-process model of fly sleep. Here we describe the adaptation of such a model, identify the sleep homeostatic parameters that best explain fly sleep, and use it to reveal interesting relationships between sleep and the circadian clock and highlight the utility of this quantitative model in understanding fly sleep. Based on this work, we also propose that longer durations of inactivity, which have been recognized by others as reflecting a relatively deep sleep state, are likely a stronger reflection of sleep homeostatic processes in *Drosophila* than the currently employed unitary definition of fly sleep as any bout of inactivity lasting five minutes or more.

## Results

### Circadian time-series analysis of *Drosophila* sleep

Though sleep is considered a major output of the circadian system, sleep behavior in flies has rarely been subjected to circadian time-series analysis. The vast majority of studies use locomotor activity, typically measured as infrared beam crossings, for such analyses. However, to fully understand the circadian control of sleep, time-series analyses should be performed directly on sleep data. For this reason, we have developed an analysis package [40], which is now available on the Comprehensive R Archive Network (CRAN; https://cran.r-project.org/web/packages/phase/index.html). Like double plotted actograms for locomotor as activity, sleep rhythms can be visualized as double plotted “somnograms” (Fig. 1B). Consistent with previous work [12,14,30], somnograms reveal two distinct phases of increased sleep under light/dark (LD) cycles, one during the daytime and one during the nighttime (Fig. 1B). Although the field has long recognized that fly sleep is bimodal, in that there are two major episodes of sleep in each diurnal or circadian cycle [12,14,30], traditional time-series methods to quantitatively assess bimodality and periodicity in sleep time-series have not been employed. We used the Chi-squared (*χ^2^*) periodogram [41] on sleep time-series data to analyze the two daily peaks of sleep under both LD cycles and constant darkness (DD), revealing two clear peaks in the periodogram, one at 12-h and the other at 24-h (Fig. 1C). Though largely similar, there are subtle differences between the periodograms of sleep and activity data measured under LD cycles and DD (Figs. 1C and 1D, compare across panels and see Table 1). The powers of average sleep time-series are higher than the powers of locomotor activity time-series (see Table 1). This is not a mere consequence of using the Chi-square periodogram, whose limitations for the quantitative assessment of rhythmic power has been described [42–44], as similar results were also obtained using the Lomb-Scargle periodogram (Suppl. Table 1).

**Table 1:**
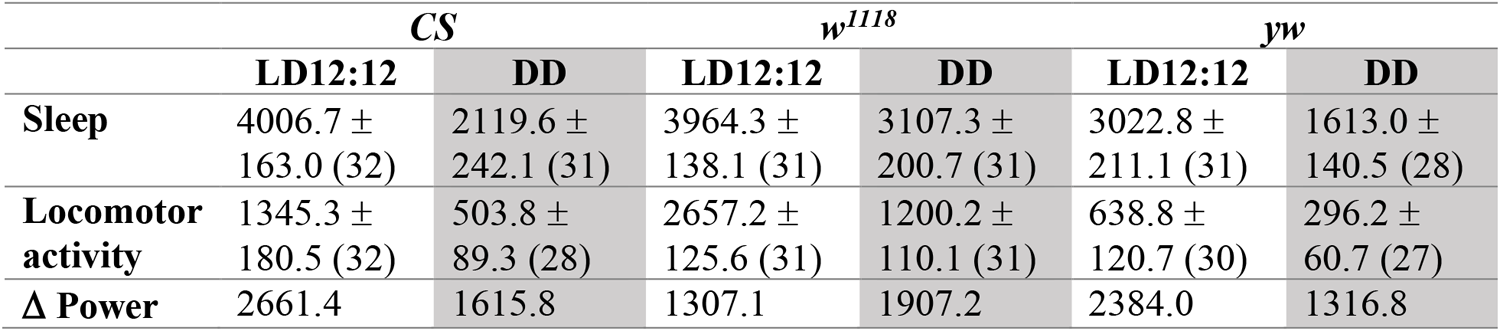
Adjusted power values (mean±SEM) of *Canton-S* (CS), *w^1118^*and *yw* flies from Chi-square periodogram analyses on sleep and locomotor activity time-series under LD and DD conditions in the circadian range (between 16- to 32-h). Also reported are the differences in power values between sleep and locomotor activity powers for each genotype. Sample sizes are provided in parentheses.

Using the Chi-square periodogram, the 12-h peak is typically lower than the 24-h peak in periodograms, but this difference is much smaller for the locomotor activity periodograms compared to sleep periodograms – in other words, the 12- and 24-h periodogram peaks are of more comparable height for locomotor activity (compare Figs. 1C and 1D). Therefore, we emphasize that irrespective of the periodogram method used, it is critical to employ sleep data to perform long-term time-series analyses rather than using locomotor activity, if one is to address the circadian regulation of sleep directly. Next, using the last four cycles under entrainment, we estimated the percentage of a 24-h day that a fly spends sleeping. Across the three genotypes, we found that flies spend ∼70 to 80% of their day sleeping (Fig. 1E). We used these two general features of sleep under LD conditions, the presence of two daily peaks in the periodogram (at 12- and 24-h under LD) with approximately 75% of time spent sleeping within a 24-h cycle, to develop our two-process model of fly sleep.

### Deriving the *Drosophila* two-process model

Although the original two-process model described above (Fig. 1A) produces a single daily peak of sleep, some animals, including humans, often display biphasic/bi-episodic sleep with a nap during mid-day and deeper sleep during the night, much like sleep in *Drosophila* [15,45]. S- and C-process parameters that could produce bimodal sleep (e.g., daytime, and nighttime sleep episodes) were identified in an earlier two-process model [5]. We therefore began building our model with previously published parameter values defining the C-process that were shown to produce bimodal sleep and created various models consisting of different combinations of build-up and decay time-constants of sleep pressure (see Suppl. Fig. 1). It is imperative to explicitly state here that throughout this study we operate under the assumption that the sleep pressure build-up and decay rates themselves are constant over time, which may not be the case in the real world. S-process parameters were determined in humans using EEG-based measures of sleep states [5]. In the absence of recognized physiological correlates of fly sleep, the use of behavioral inactivity was the only method by which we could estimate homeostatic parameters in this system. In principle, electrophysiological correlates are a proxy for sleep depth [31], and so are behavioral correlates. Therefore, we use behavioral inactivity states and estimate S-process parameters. For each of these models we first asked if they produced mono-episodic, bi-episodic, pluri-episodic, or arrhythmic/unstructured sleep patterns (see Suppl. Fig. 1). We found that a large region of the parameter space of S-process (all values that S can have, as defined by the input to the model) produced pluri-episodic sleep rhythms (Fig. 2A), meaning they had three or more peaks above the significance line in a periodogram analysis and had three or more episodes of sleep per cycle. A significant region of the parameter space failed to produce coherent sleep rhythms (Fig. 2A), meaning that there were no detectable periodicities revealed by our *χ^2^* analysis. Smaller regions of the parameter space produced coherent mono-episodic (exactly one sleep episode/24-h) or bi-episodic (exactly two sleep episodes/24-h) sleep rhythms (Fig. 2A).

**Figure 2:**
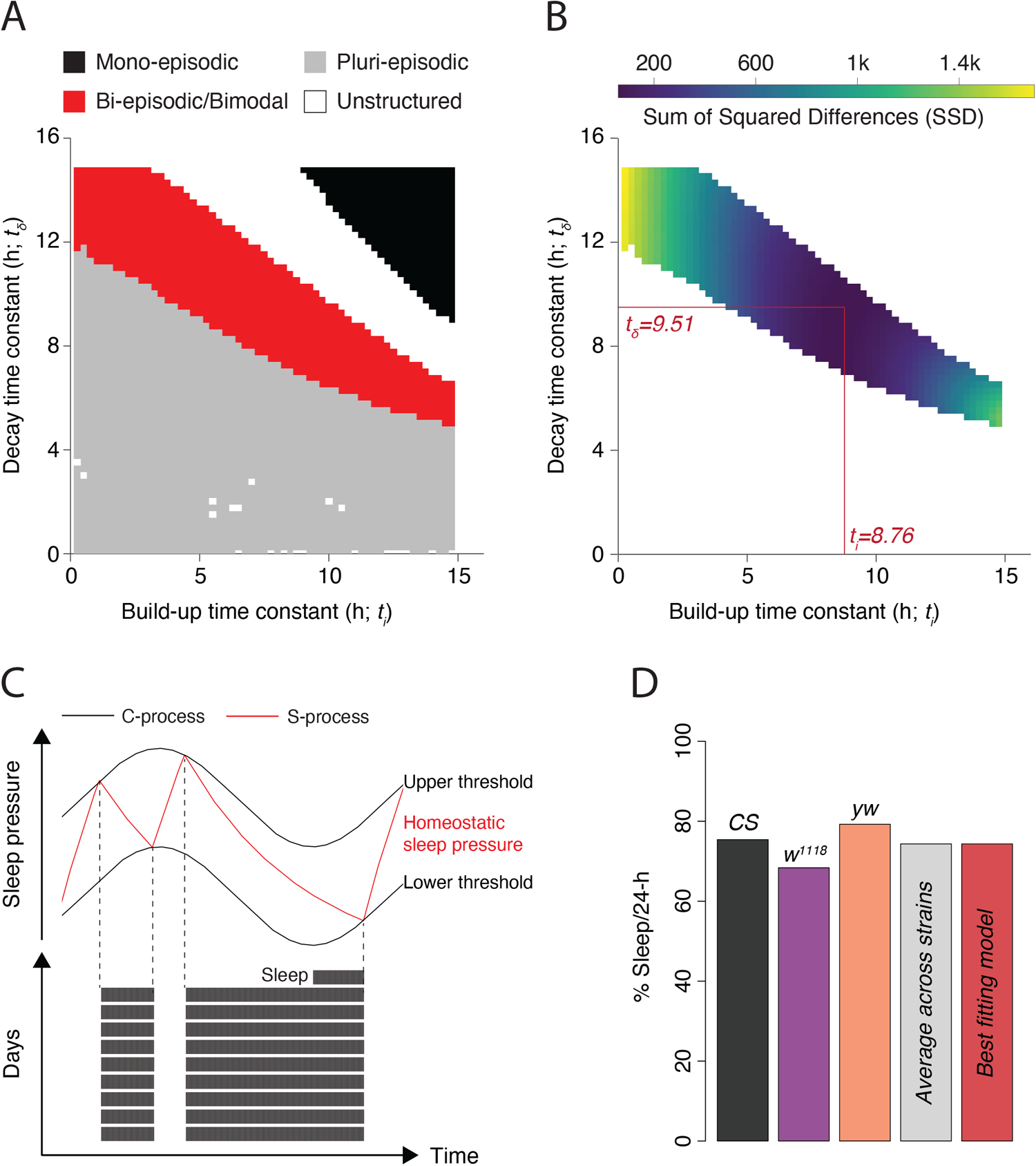
Building a *Drosophila* two-process model of sleep regulation. (A) Heatmap summarizing the nature of sleep rhythms within the build-up and decay time constant parameter space for sleep pressure. (B) Heatmap showing the sum of squared differences (SSD) between model prediction and empirical values in the same parameter space for the value combinations producing bi-episodic/bimodal rhythms. The red lines along the *x*- and *y*-axes represent the sleep pressure parameters that best fit empirical data. (C) The two-process model with best-fitting parameters describing fly sleep. (D) The percentage of time spent sleeping/24-cycle in the three genotypes and their average value compared with the predicted value from the model.

Given that we were interested in parameter values that produce two daily episodes of sleep, we subsequently focused on combinations of time constants that yielded exactly two episodes of sleep/24-h cycle. Among these, we quantified the percentage of time spent sleeping per cycle. For each set of build-up and decay parameters, we then compared the values they produced with the percent of time that three fly genotypes, *Canton-S*, *w^1118^* and *yw*, slept per cycle (median over all flies/genotype), using sum of squared differences (SSD; see Materials and Methods; Fig. 2B). We found that the lowest SSD (best fit) was attained when the value for a build-up time constant of 8.76-h and a decay time constant of 9.51-h (Fig. 2B). Therefore, we chose these parameter values as the best descriptors for daily fly sleep under standard LD cycles. The model generated using these parameter values is shown in Fig. 2C. Though it appears to underestimate daytime sleep and somewhat overestimate nighttime sleep, the percentage time spent sleeping/24-h cycle predicted by the model was ∼74.36%, which is close to the percentage of time that the three fly strains spend asleep individually and identical to the average value across the three strains (Fig. 2D). It is curious that the build-up time constant in flies is faster than the decay time constant. This appears to contrast with rodent systems and humans wherein the decay time constant is much faster than the build-up [46]. We think that this could reflect the relative amounts of sleep and wake within a 24-h cycle. Rodents and humans are awake longer than they are asleep, which is the opposite for flies. The short burst of locomotor activity in flies may impose constraints on the rates of sleep pressure changes.

### Predictions from the two-process model

We built a two-process model of *Drosophila* sleep to systematically predict the effects of altering either the S- or the C-process on fly sleep rhythms. We first asked what would happen to daily sleep if the circadian modulation of sleep thresholds were abolished. To model this, we set the amplitude of the upper and lower thresholds to zero, such that they were held constant at their mean values. Our model made the straightforward prediction that eliminating the contribution of the circadian system would result in ultradian sleep rhythms (Figs. 3A and B). Ultradian rhythms in the locomotor activity of flies with no functional circadian clocks have been reported previously [47,48]. Our model suggests that such rhythms are a predictable consequence of a functional S-process in the absence of circadian modulation of sleep thresholds and predicts ultradian periodicities of sleep of ∼10-h in the absence of a functional circadian clock.

**Figure 3:**
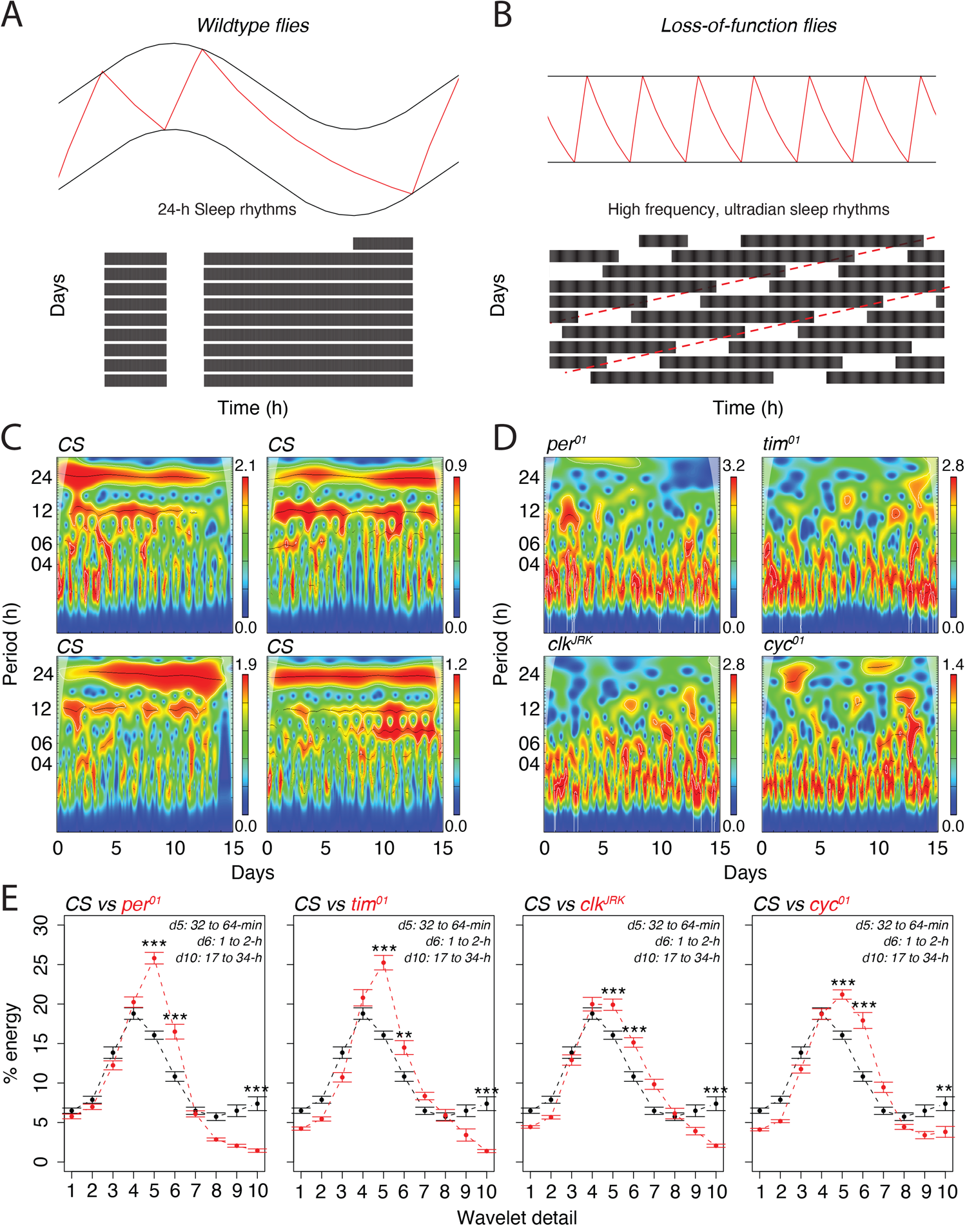
Ultradian periodicities are a predictable consequence of a functional S-process under a dysfunctional C-process. (A) The wildtype two-process model (top) and the somnogram of simulated sleep data produced by this model (bottom). (B) The same model as in panel (A) but with flat, non-oscillating thresholds to model sleep regulation in a loss-of-function clock mutant (top) and the somnogram it produced (bottom). (C) Four representative continuous wavelet transform (CWT) scalograms for *Canton-S* (CS; C) and (D) one each for *per^01^*, *tim^01^*, *clk^JRK^* and *cyc^01^*. The color bars against each scalogram are representative of local rhythmic power. Note: the *y*-axes on the scalograms are on a logarithmic scale, and the data are binned in 60-min intervals. (E) Wavelet detail-specific characterization of sleep/wake rhythms (binned in 1-min intervals) show the % energy at each scale. Details 5 and 6 correspond to ultradian periods ranging from 32-min to 2-h. Detail 10 corresponds to the circadian range (17- to 34-h). The wildtype data set is replotted in each panel to facilitate comparisons with the four mutant lines. Values plotted are mean±SEM. See text for details. For each detail level, two-sample t-tests were carried out for comparing wildtype and mutant flies, which would result in 10 overall comparisons. The *p*-values reported were adjusted for multiple comparisons, using a Benjamini-Hochberg correction. Statistical results are shown only for details 5 (*per^01^ vs CS*: *t56.02* = –10.85, *p* = 2.15e-14; *tim^01^ vs CS*: *t49.07* = –8.69, *p* = 1.69e-10; *clk^JRK^ vs CS*: *t56.66* = –4.36, *p* = 0.0001; *cyc^01^ vs CS*: *t61.04* = –6.53, *p* = 1.45e-07), 6 (*per^01^ vs CS*: *t53.14* = –5.08, *p* = 9.87e-06; *tim^01^ vs CS*: *t55.05* = –3.42, *p* = 0.002; *clk^JRK^ vs CS*: *t61.98* = –5.06, *p* = 1.35e-05; *cyc^01^ vs CS*: *t50.95* = –6.03, *p* = 6.12e-07) and 10 (*per^01^ vs CS*: *t34.95* = 6.58, *p* = 6.76e-07; *tim^01^ vs CS*: *t34.12* = 6.68, *p* = 5.57e-07; *clk^JRK^ vs CS*: *t34.27* = 5.92, *p* = 7.66e-06; *cyc^01^ vs CS*: *t58.66* = 3.21, *p* = 0.003). * < 0.05, ** < 0.01, *** < 0.001, **** < 0.0001, and N.S. (Not Significant).

We first visualized this prediction by examining sleep rhythms in wildtype and various loss-of-function clock mutants (*per^01^, tim^01^, clk^JRK^ and cyc^01^*) under constant darkness using continuous wavelet transforms (CWT; a method to detect wide range of periodicities at local timescales) as have been used previously for circadian time-series data (see Materials and Methods) [49–52]. We then used maximum overlap discrete wavelet transforms (mo-DWT) to examine the extent of variance explained by specific ranges of period values for each of these strains as has been done before [53].

A close inspection of the scalograms in Fig. 3C show the clear presence of strong circadian rhythms (the red band around 24-h) of sleep, in wildtype CS flies. Interestingly, clear ultradian sleep rhythms are also visible in these flies. On the other hand, representative scalograms for four loss-of-function clock mutants show no presence of circadian rhythms in sleep, as expected. These flies show significantly stronger ultradian rhythms in sleep (Fig. 3D) as compared to their wildtype counterparts. We then used the mo-DWT to quantitatively characterize the scale-based percentage energy associated with each level of wavelet detail. A wavelet detail is associated with a specific range of periodicities that is defined by *2^j^Δt* to *2^j+1^Δt*, where *j* is the wavelet detail level and *Δt* is the sampling interval or data collection bin size. Therefore, for data collected in 1-min bins, wavelet detail level 5 would correspond to periodicities ranging from *2^5^×1* = 32-min to *2^6^×1* = 64-min. When we performed such characterization, we found that there was consolidation of energy in wavelet details 3-6 and 10 in the wildtype flies. Given that our data was sampled every minute, wavelet details 3-6 are associated with periods ranging from ∼8-min to ∼2-h. Wavelet detail 10, as described above, is the circadian band with periods ranging from 17- to 34-h (*2^10^×1* = 1024-min

= 17.067-h to *2^11^×1* = 2048-min = 34.13-h). We found that wildtype flies had significantly higher percentage of energy associated with wavelet detail 10 – the scale index associated with circadian periodicities – compared to all four loss-of-function clock mutants (Fig. 3E). Importantly, in all four clock mutants, there was significantly higher energy in wavelet details 5 and 6 (corresponding to ∼0.5- to ∼2-h periods) compared to wildtype flies (Fig. 3E). It is also interesting that there was a broad distribution of ultradian periodicities ranging predominantly from ∼16-min to ∼2-h (Figs. 3C-E) in these flies that were apparent in raw sleep time-series (example shown for *per^01^* flies in Suppl. Fig. 2). The range of ultradian periodicities that we detected for sleep was similar to those detected previously for locomotor activity [47,48]. Our results indicate that sleep/wakefulness rhythms are indeed ultradian in loss-of-function clock mutants and are likely driven by a functional S-process in the absence of circadian clock-controlled thresholds, much like a relaxation oscillator. We think that this provides a more parsimonious explanation for the presence of ultradian rhythms in activity than the uncoupling of endogenous ultradian oscillators in loss of function clock mutants [47,48,54].

### How independent are the S- and C-processes in *Drosophila*?

Based on the persistence of the C-process in the absence of sleep and the maintenance of the S-process in the absence of circadian timekeeping, the two-process model assumes that they are produced by distinct mechanisms. However, it is not clear whether these separate processes operate independently, i.e., without influencing one another’s kinetics (reviewed in [11]). Rhythms in the propensity to sleep appear to be independent of the duration of prior waking or sleep duration, implying the independent action of the clock and the homeostat [31]. Furthermore, homeostatic increases in NREM sleep and slow wave activity are observed after sleep deprivation in the absence of the brain’s central circadian pacemaker, the suprachiasmatic nuclei (SCN) [55–58]. Though these results support the separateness of the S- and C-processes, recent studies have suggested that these two processes influence one another (reviewed in [11]). For example, loss-of-function modifications to the circadian clock appear to have significant impact on sleep homeostatic markers in mice and flies [32,59–63]. Furthermore, variation in human chronotypes is associated with variation in sleep homeostatic markers (reviewed in [64]). How might the separate S and C processes influence one another in *Drosophila*?

Thanks to the genetic tools available for the manipulation of the C-process in *Drosophila*, we were able to generate flies with a wide range of circadian periodicities and assess the relationships between the period of the C-process and the amount and waveform of sleep. Thus, we can compare the model’s predictions to the experimental effects of C-process manipulation on sleep rhythms (Fig. 4A). First, we recorded locomotor activity in flies with a wide range of circadian periods under DD and constant temperature conditions. We employed five *period* alleles (located on the X-chromosome): *per^T^* (free-running period – 15-h; [65]), *per^S^* (free-running period – 21-h; [66]), *per^+^* (free-running period – 23.5-h), *per^L^* (free-running period – 27.75-h; [66]) and *per^01^* (arrhythmic; [66]). As expected, these alleles produce significantly different free-running periods (see Table 2). By creating all possible combinations of these alleles (by including heterozygous females), we generated large numbers of flies with a continuous and wide range of free-running circadian periods between ∼15- to ∼30-h, as previously described by others (see Table 2; [65,66]). We performed this experiment twice, pooled flies from both experiments, and quantified power and hours of sleep/circadian cycle for all rhythmic flies observed (Fig. 4B-C; Tables 2 and 3).

**Figure 4:**
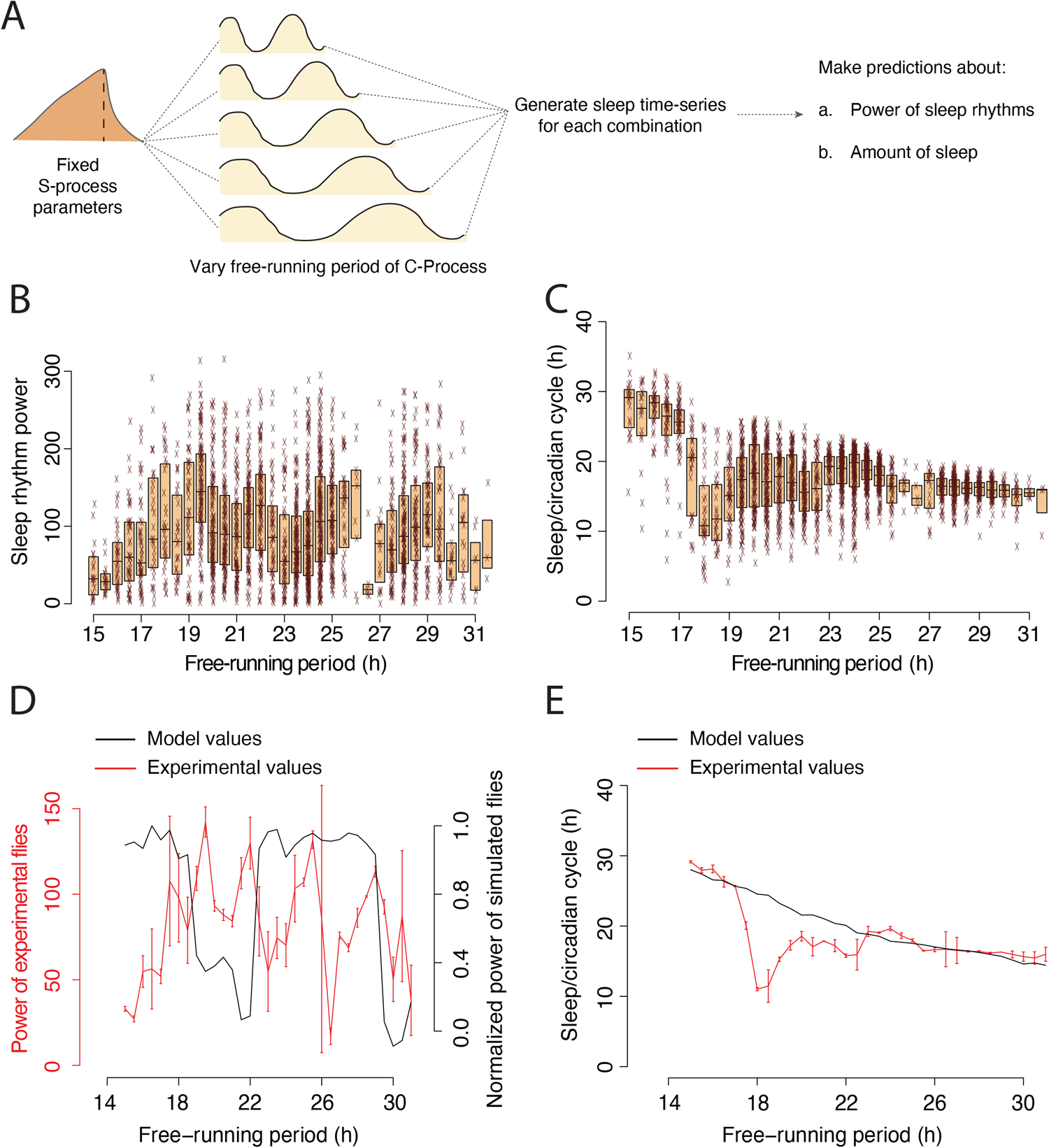
The model predicts regions of relatively weak sleep rhythms across a range of free-running periods which do not match empirical data. **(A)** Schematic representation of the simulation workflow upon which predictions were made. (B) Power of sleep rhythms and (C) time spent sleeping/circadian cycle for flies with different free-running periods, generated through combinations of five *period* alleles (see Materials and Methods). (D) Median power of sleep rhythms (mean±SEM across two replicate runs) and (E) time spent sleeping/circadian cycle (mean±SEM across two replicate runs) across the entire range of free-running periods overlayed with the predictions of the two-process model when only the free-running period was altered in the model.

**Table 2:**
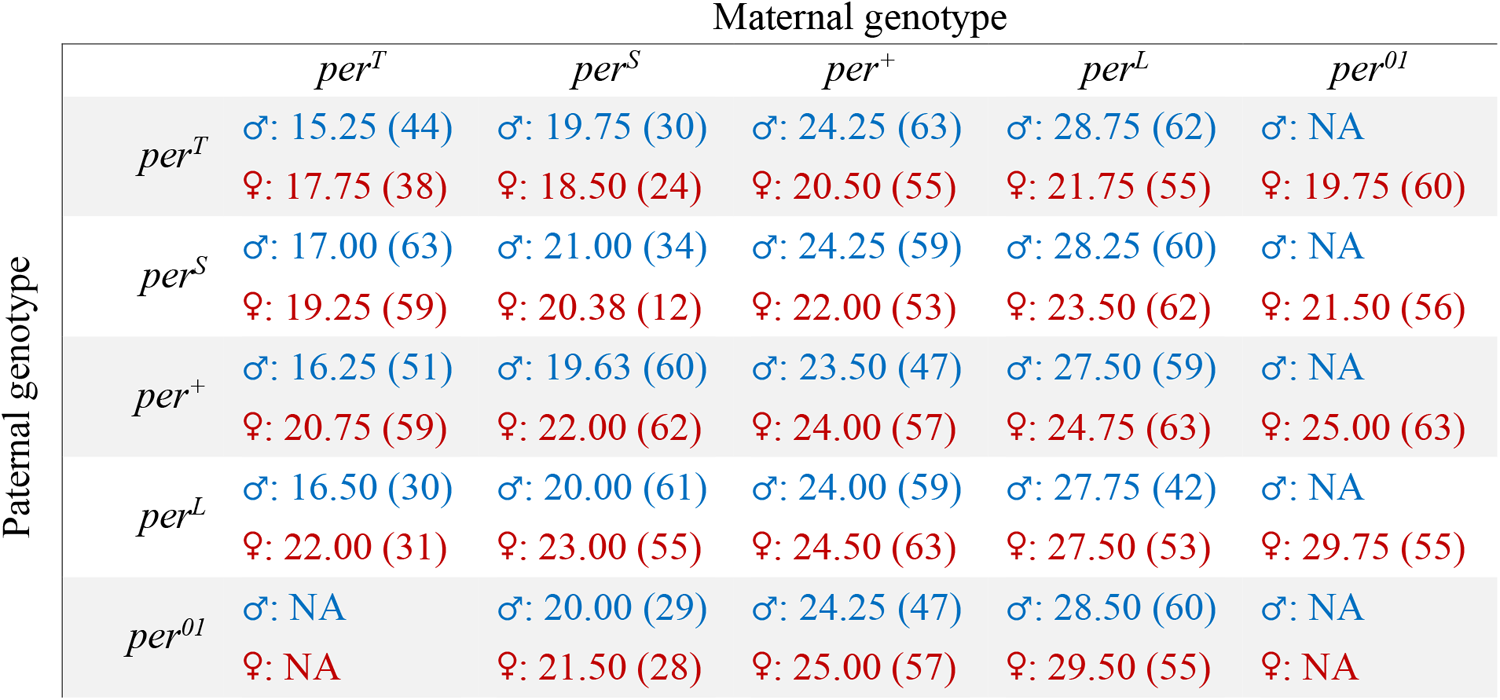
Median free-running periods of sleep rhythms under constant darkness averaged across two replicate runs. The values for male (top, blue) and female (bottom, red) progeny from each cross are reported. Outcomes of any cross that had fewer than 12 flies pooled from both replicate runs have not been reported in this table. Sample sizes are provided in parentheses.

|  |  | Maternal genotype |  |  |  |  |
| --- | --- | --- | --- | --- | --- | --- |
|  |  | *per<sup>T</sup>* | *per<sup>S</sup>* | *per<sup>+</sup>* | *per<sup>L</sup>* | *per<sup>01</sup>* |
| Paternal genotype | *per<sup>T</sup>* | ♂: 15.25 (44) | ♂: 19.75 (30) | ♂: 24.25 (63) | ♂: 28.75 (62) | ♂: NA |
|  |  | ♀: 17.75 (38) | ♀: 18.50 (24) | ♀: 20.50 (55) | ♀: 21.75 (55) | ♀: 19.75 (60) |
|  | *per<sup>S</sup>* | ♂: 17.00 (63) | ♂: 21.00 (34) | ♂: 24.25 (59) | ♂: 28.25 (60) | ♂: NA |
|  |  | ♀: 19.25 (59) | ♀: 20.38 (12) | ♀: 22.00 (53) | ♀: 23.50 (62) | ♀: 21.50 (56) |
|  | *per<sup>+</sup>* | ♂: 16.25 (51) | ♂: 19.63 (60) | ♂: 23.50 (47) | ♂: 27.50 (59) | ♂: NA |
|  |  | ♀: 20.75 (59) | ♀: 22.00 (62) | ♀: 24.00 (57) | ♀: 24.75 (63) | ♀: 25.00 (63) |
|  | *per<sup>L</sup>* | ♂: 16.50 (30) | ♂: 20.00 (61) | ♂: 24.00 (59) | ♂: 27.75 (42) | ♂: NA |
|  |  | ♀: 22.00 (31) | ♀: 23.00 (55) | ♀: 24.50 (63) | ♀: 27.50 (53) | ♀: 29.75 (55) |
|  | *per<sup>01</sup>* | ♂: NA | ♂: 20.00 (29) | ♂: 24.25 (47) | ♂: 28.50 (60) | ♂: NA |
|  |  | ♀: NA | ♀: 21.50 (28) | ♀: 25.00 (57) | ♀: 29.50 (55) | ♀: NA |

**Table 3:** Median Chi-square periodogram power of sleep rhythms under constant darkness averaged across two replicate runs. The values for male (top, blue) and female (bottom, red) progeny from each cross are reported. Outcomes of any cross that had fewer than 12 flies pooled from the replicate runs have not been reported in this table. Sample sizes are provided in parentheses.

|  |  | Maternal genotype |  |  |  |  |
| --- | --- | --- | --- | --- | --- | --- |
|  |  | *per<sup>T</sup>* | *per<sup>S</sup>* | *per<sup>+</sup>* | *per<sup>L</sup>* | *per<sup>01</sup>* |
| Paternal genotype | *per<sup>T</sup>* | ♂: 33.0 (44) | ♂: 187.5 (30) | ♂: 136.1 (63) | ♂: 135.5 (62) | ♂: NA |
|  |  | ♀: 33.6 (38) | ♀: 143.9 (24) | ♀: 79.5 (55) | ♀: 98.4 (55) | ♀: 119.4 (60) |
|  | *per<sup>S</sup>* | ♂: 98.0 (63) | ♂: 109.7 (34) | ♂: 95.9 (59) | ♂: 137.5 (60) | ♂: NA |
|  |  | ♀: 110.7 (59) | ♀: 40.7 (12) | ♀: 105.1 (53) | ♀: 70.8 (62) | ♀: 103.3 (56) |
|  | *per<sup>+</sup>* | ♂: 51.5 (51) | ♂: 161.0 (60) | ♂: 79.4 (47) | ♂: 64.9 (59) | ♂: NA |
|  |  | ♀: 76.4 (59) | ♀: 117.0 (62) | ♀: 92.6 (57) | ♀: 118.0 (63) | ♀: 130.3 (63) |
|  | *per<sup>L</sup>* | ♂: 93.7 (30) | ♂: 81.1 (61) | ♂: 53.9 (59) | ♂: 72.0 (42) | ♂: NA |
|  |  | ♀: 108.4 (31) | ♀: 55.5 (55) | ♀: 134.0 (63) | ♀: 85.4 (53) | ♀: 92.8 (55) |
|  | *per<sup>01</sup>* | ♂: NA | ♂: 114.6 (29) | ♂: 32.7 (47) | ♂: 56.1 (60) | ♂: NA |
|  |  | ♀: NA | ♀: 114.4 (28) | ♀: 102.4 (57) | ♀: 72.7 (55) | ♀: NA |

We also simulated sleep rhythms by altering the free-running periods of the upper and lower thresholds in our model (see Materials and Methods), leaving all other parameters unchanged (Fig. 4A; Suppl. Fig. 3); we used these simulations to quantify the power of sleep rhythms and the amount of sleep/circadian cycle. Our model predicted that specific ranges of free-running periodicities would be associated with relatively weak sleep rhythms (Fig. 4D). Specifically, that sleep rhythms would display relatively low periodogram powers for circadian periods between 18- and 22-h, and beyond ∼29-h. The model also predicted relatively higher power for periods between 15- to 18-h and 22- to 29-h (Fig. 4D). Though the power of sleep rhythms in our experimental flies did vary markedly across circadian periods, our model’s predictions did not match experimental results, which revealed that sleep rhythm power displays considerable variation across the entire range of periodicities examined, with several troughs displayed across the ranges of periods observed (Fig. 4D). When we examined the amount of sleep/circadian cycle in our simulations, we found a monotonic decline in the hours of sleep with lengthening circadian period (Fig. 4E). Once again, our experimental results differed from our model’s predictions, with experimental flies displaying lower amounts of sleep than predicted between ∼18- and 20-h (Fig. 4E).

We also quantified the relationship between free-running period and the power of locomotor activity rhythms. Interestingly, we found that there is a reasonable match in normalized power of locomotor activity and sleep rhythms in the short period range (between ∼15- to ∼21-h), but there is quite a mismatch for period values beyond ∼21-h (Suppl. Fig. 4A). Consequently, the degree of match between power of locomotor activity time-series and predictions from the model is different from that with sleep time-series (compare Figs. 4D and Suppl. Fig. 4B), thereby highlighting the importance of using sleep time-series to study the circadian regulation of sleep rather than extrapolate from periodogram analyses on locomotor activity time-series. Furthermore, this quantitative relationship between free-running period and power of sleep rhythms was not a consequence of the limitations of the Chi-square periodogram. Analysis using the Lomb-Scargle periodogram revealed a similar relationship between period and power (Suppl. Fig. 5).

Although the S- and C-processes were initially thought to act without influencing each other’s kinetics [31], recent work indicates that they do in fact influence each other [11,32,59–64]. Our model, which assumes that these two processes function independently, failed to precisely predict the relationship between clock speed and sleep rhythm power and amounts of sleep. We therefore sought to adjust the S-process parameters for each free-running period to maximize the fit with our experimental data, once again using the SSD-based approach (Fig. 5A; also see Suppl. Fig. 6). We did this by exploring the build-up and decay rate parameter space for each fly and identified the best fitting set, thereby producing a much better agreement for power and amount sleep between the model and experimental results (see Materials and Methods; Figs. 5B and 5C). The S-process values that produce this fit are shown in Fig. 5D. We found that, there was no linear relationship between free-running period and the build-up time constant of sleep pressure (Fig. 5D-left). On the other hand, we found a statistically significant linear relationship between free-running period and sleep pressure decay time constant, wherein flies with longer free-running periods (flies with slower clocks) also have higher time-constants of sleep pressure release (slower decay of sleep pressure; Fig. 5D-right). Therefore, our results show that the rates of the S- and C-processes are indeed interdependent in a way that has implications for our understanding of the regulation of *Drosophila* sleep.

**Figure 5:**
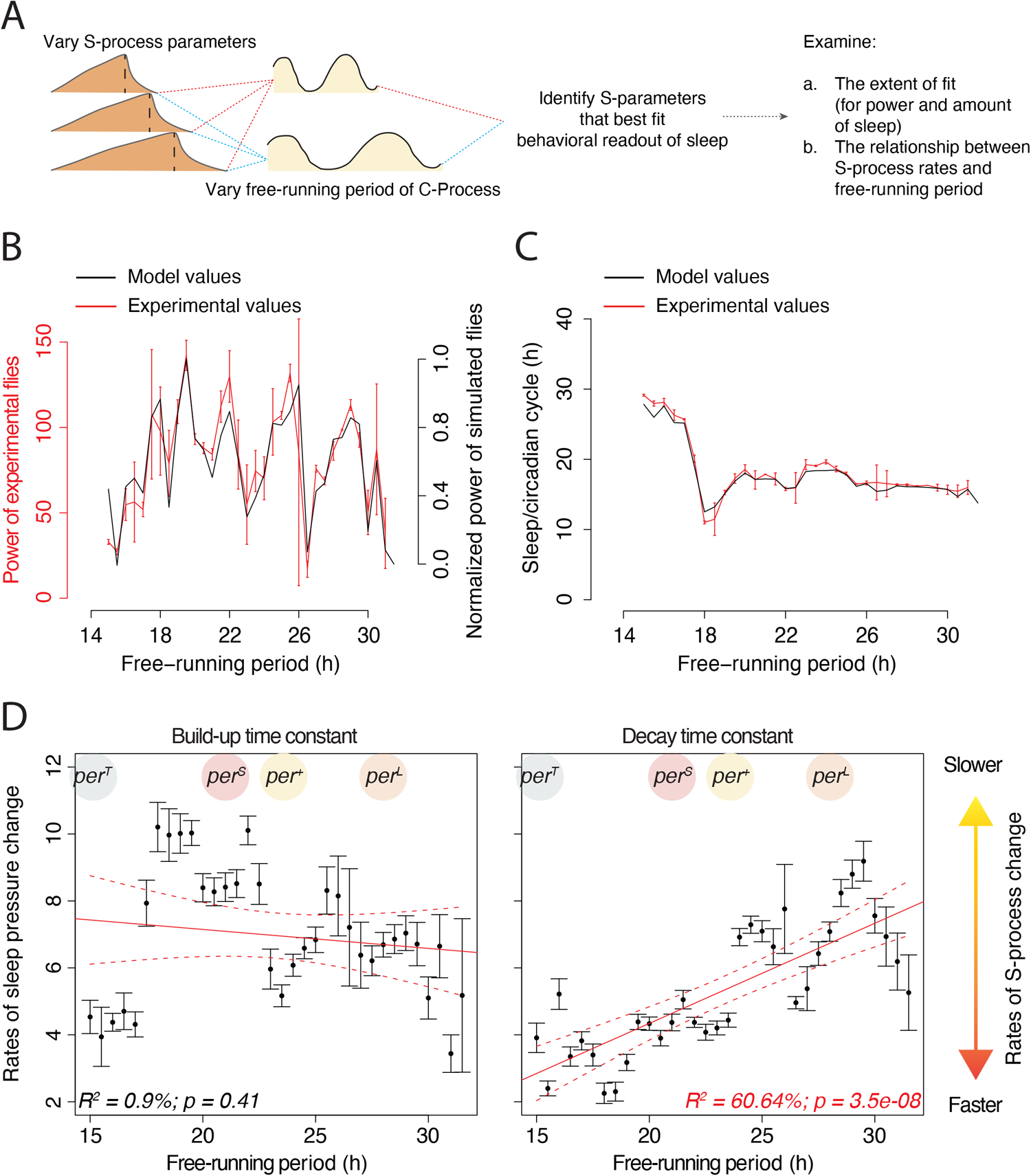
Allowing homeostatic parameters to change with free-running periods suggest a strong inter-dependence between the S- and C-processes. (A) Schematic representation of the simulation routine to estimate best fitting S-process parameters for each individual fly. (B) Median power of sleep rhythms (mean±SEM across two replicate runs) and (C) time spent sleeping/circadian cycle (mean±SEM across two replicate runs) across the entire range of free-running periods overlayed with the predictions of the two-process model when build-up and decay time constants were allowed to covary with the free-running period. Note that the experimental data shown in panels (B) and (C) are the same as those shown in Figs. 4D and 4E and have been replotted here to facilitate comparisons. (D) The relationship between build-up time constant (left) and decay time constant (right) of sleep pressure and free-running periods. Values plotted are mean±SEM. The red solid line is the estimate from a linear regression model and the dotted red lines are the 95% confidence intervals. Also shown are the regions along the *x*-axis corresponding to the free-running periods of males bearing the four rhythmic *period* alleles used in our study (see also Table 2).

### To what extent is *Drosophila* sleep a unitary state?

Although sleep has largely been treated as a unitary state in the fly, several independent approaches support the notion that long durations of inactivity correspond to a relatively “deep-sleep” stage [33–39]. For example, metabolic rates of flies that have been inactive for ∼35-min are significantly lower than flies that have been inactive for the standard five-minute definition of fly sleep [33]. Furthermore, experiments tracking arousal thresholds and brain physiology are consistent with deeper sleep intensities when flies have been inactive for ∼30-min [34].

The original two-process model made predictions regarding the amount of total sleep time by modeling the S-process based on the exponential decay of the power of sleep EEG (0.75- to 25-Hz). This wide range of EEG frequencies corresponds to all sleep types, i.e., slow wave and REM sleep. Interestingly, the decay constant derived from such a wide range of EEG frequencies was very similar to the constant derived from slow wave sleep alone (0.5- to 2-Hz) [5,67], thereby suggesting that although the two-process model accounts for total sleep, the S-process properties are most closely reflected by NREM sleep. Since decay rate constants best reflect NREM sleep, we assessed the degree to which our model predictions would describe empirical data when increasing durations of inactivity were used as the defining criterion for a sleep like state, thereby biasing our sleep analysis to longer, presumably deeper, bouts of sleep in experimental flies.

Because recent studies in flies (discussed above) have established a clear correlation between the duration of inactivity and the depth of sleep, we use the term “deep” to refer to sleep bouts that are longer than the traditional unitary five-minute inactivity definition of fly sleep. Similar associations between sleep consolidation (number of spontaneous brief awakenings) and sleep pressure (measured using slow wave activity) have also been reported in rodent models [68], therefore suggesting that this association is likely a conserved phenomenon.

We systematically altered the inactivity duration definition of sleep, i.e., the duration of inactivity used to define the sleep state and examined the correlation between the amount of time flies spent sleeping and our model’s prediction (Figs. 6A-B). Traditionally, any bout of inactivity that is five minutes or more is defined, in flies, as sleep. The correlation between our model’s prediction and measurements of amount of time spent sleeping in our series of allelic combinations of *period* changes very little with increasing inactivity definitions, at least until the criterion for sleep reaches 30-min of inactivity (Figs. 6A-B). This result implies that shorter bouts of inactivity contribute very little to how the amount of time spent sleeping varies with circadian period.

**Figure 6:**
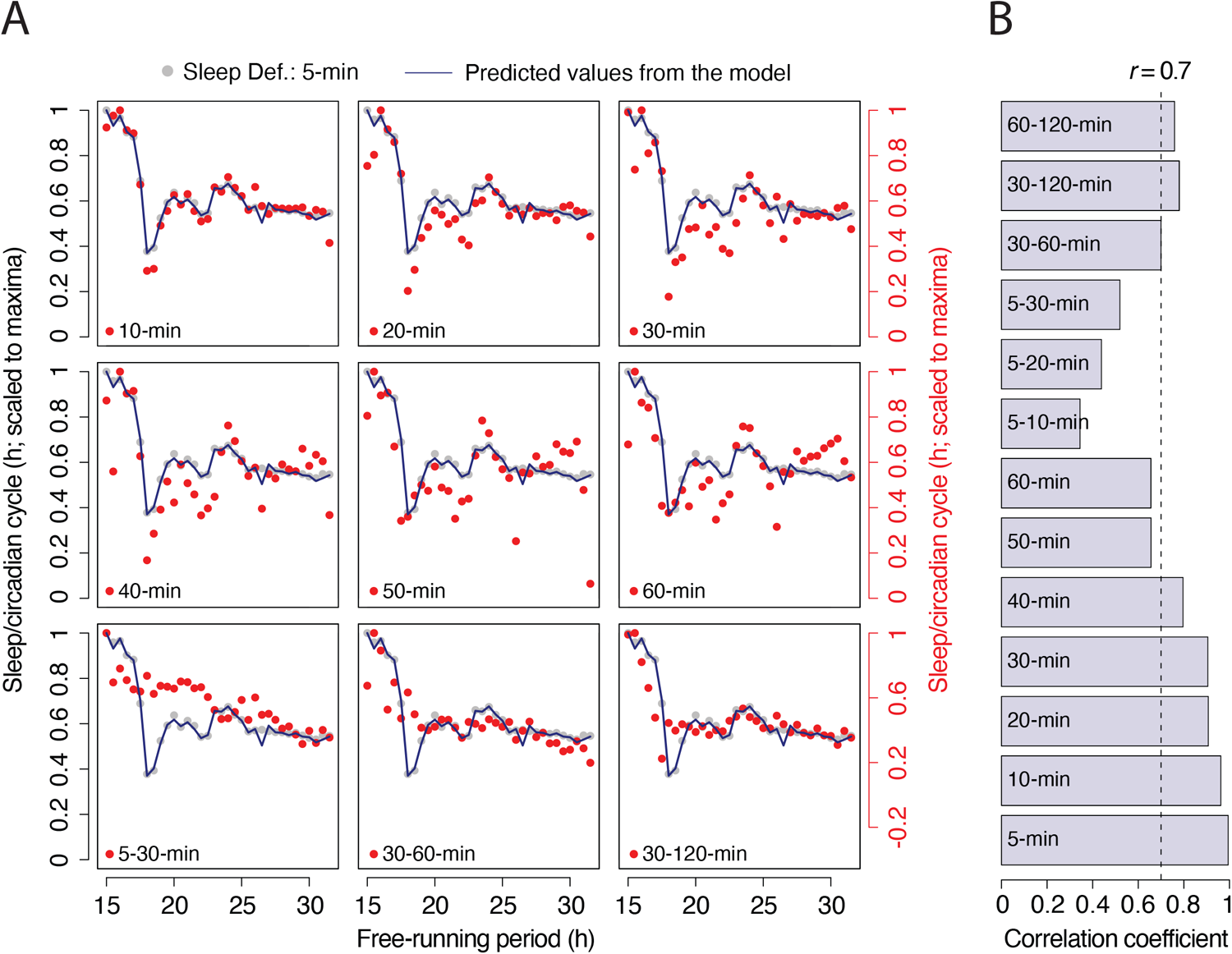
Short bout sleep is dispensable for capturing the relationship between circadian period and amount of sleep. (A) Scatter plots of the amount of time spent sleeping/circadian minute across different free-running periods for different inactivity duration-based definitions of sleep overlayed with the amounts of sleep measured using standard definition of sleep and the amounts predicted by the model. All the sleep amounts are scaled to maxima so as to restrict the *y*-axis between 0 and 1. Note that the same model and standard definition of sleep data are re-plotted in every panel to facilitate comparisons. For each definition of sleep (shown in red), the *y*-axis is plotted on the right side of the plot. In case of the bottom three panels, the minima values on the secondary *y*-axis (in red) have been adjusted to enable easy visual comparisons. Note that data for the amount of sleep using the 5-min definition of sleep are the same as that depicted in Figs. 4E and 5C. Each dot in all the panels represents the median value of flies within each detected free-running period, averaged over two independent replicate runs. (B) The correlation coefficients from Pearson’s product-moment correlations on the raw amounts of sleep/circadian cycle between model predictions and different inactivity duration-based definitions of sleep. The scaling of data and adjustment of the *y*-axis have no bearing on the correlation coefficients reported.

To further test this idea, we performed the same analysis as above but with specific ranges of inactivity durations used for the definition of sleep (see Materials and Methods for details). For instance, a five to 30-minute interval definition of sleep would mean that any bout of inactivity that is equal to or more than five minutes but fewer than 30 minutes would be considered sleep. We found that the correlation between empirical results and model predictions is relatively low when five to 10, 20 or 30 minutes of inactivity were used as the definitions of sleep (Fig. 6B). Correlations improved a little when the inactivity definition was set to 30 to 60 minutes and became fairly high when set to 30 to 120 minutes (Fig. 6B). These results suggest that our model can account for a large part of the relationship between circadian period and amounts of deep sleep and are consistent with recent work suggesting that different durations of inactivity are unlikely to represent the same state with respect to sleep in *Drosophila* [33–39].

### An independent test confirms that faster running clocks also have altered sleep homeostatic properties

In fitting our two-process model to sleep observed over a range of circadian periods, our results indicate that, as previously suggested, the S- and C-processes influence one another. We found that changes in the decay time constant of the S-process accompanied changes in the free-running period of the C-process (Fig. 5D-right). We therefore sought to independently test this idea that flies with shorter free-running periods will also have faster decay of sleep pressure (see Fig. 5D-right). To look at this relationship more closely we examined deep sleep (30-minutes or longer of inactivity) in *per^+^* and *per^S^* flies under conditions in which the C-process had been rendered arrhythmic, a condition which reveals ultradian rhythms driven by the operation of process S alone (see Figs. 3 and 7A). Our results described above (see Fig. 5D) leads to the following predictions in the absence of circadian cycling of sleep thresholds (therefore, no circadian gating of sleep): (i) ultradian rhythms in sleep/wake will be faster in *per^s^* mutants compared to *per^+^*flies, (ii) total amount of sleep would be lower in *per^s^* mutants compared to *per^+^* flies, and (iii) number of sleep bouts will be higher in *per^S^* mutants compared to *per^+^* flies per unit time (Fig. 7A). Based on the results in the previous section (Fig. 6), we argue that examining deep sleep in these flies will perhaps be a closer reflection of process S. *Drosophila* is arrhythmic under continuous white light (LL) owing to the constitutive CRYPTOCHROME mediated degradation of TIMELESS [69–71]. Thus, LL conditions provide a convenient means of abolishing the cycling of sleep thresholds (C-process). We therefore examined sleep behavior in *per^S^*and *per^+^* (CS) flies under LL and examined longer bouts (presumably deeper, homeostatic) of sleep (see results above and Fig. 7A). As expected, over the duration of 15 days under constant white light, there was a clear absence of any circadian rhythmicity in both *per^+^* and *per^S^* flies (Fig. 7B). Employing recently described methods [53,72], we identified three specific bands of ultradian periodicities using mo-DWT (see Materials and Methods) that account for the majority of spectral variance in the sleep timeseries of *per^+^* and *per^S^* flies (Figs. 7C and 7D). These three ranges of periods were 1.07-2.1-h (*τs or τshort*), 2.1-4.27-h (*τm or τmedium*) and 4.27-8.53-h (*τl or τlong*; Figs. 7C and 7D). We analyzed the period values in these categories separately for both these strains. We found that, as predicted, *per^S^* had significantly faster ultradian periods in the *τs* category (Fig. 7E). There was no significant between-strain difference in periods in the other two period categories (Fig. 7E). Furthermore, we found statistically significantly reduced total sleep (Fig. 7F) and increased bout numbers (Fig. 7G) in *per^S^*flies compared to *per^+^* flies. Our results, therefore, support *all three* predictions from the model, thereby lending support to the notion that shorter free-running periods are indeed associated with faster rates of sleep pressure decay.

**Figure 7:**
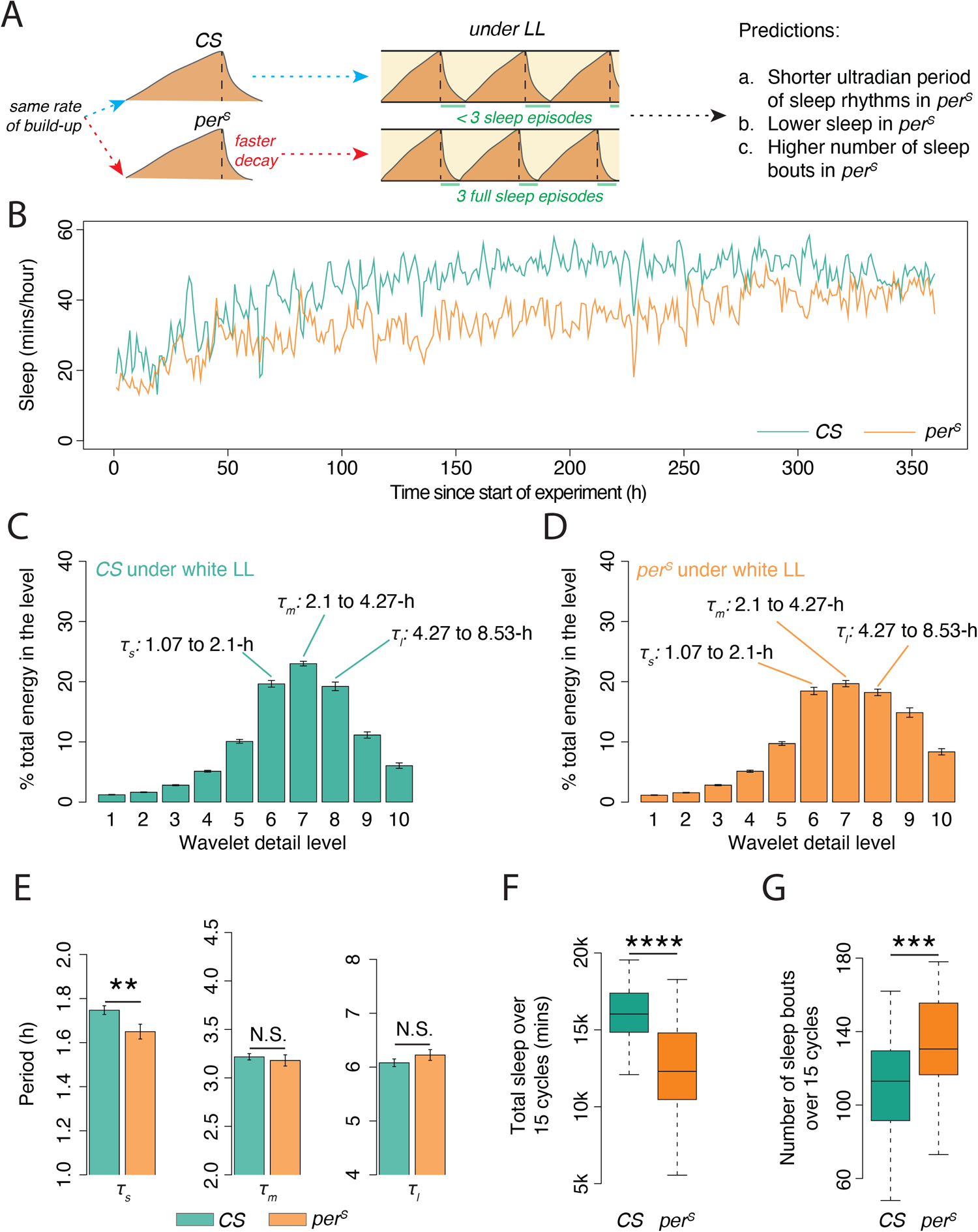
Independent tests confirm the relationship between free-running period and the rates of change of sleep homeostat. (A) Schematic representation of the predictions for short period flies having altered sleep homeostatic parameters. (B) Time-series of long bout sleep of both genotypes under continuous white light for 15 days. Discrete wavelet transform based estimate of scale-wise concentration of spectral variance in (C) CS and (D) *per^S^* flies (also see Materials and Methods). Shown are mean estimates of % energy±SEM. Note the three period bands that comprise maximal spectral energy. (E) Ultradian periods (mean±SEM) for the two genotypes under LL as estimated from the continuous wavelet transforms (Results based on one-tailed bootstrapped Welch two-sample *t*-tests – *1s*: *t30.72* = 2.49, *p* = 0.004; *1m*: *t29.46* = 0.57, *p* = 0.30; *1l*: *t35.58* = −1.19, *p* = 0.88). (F) Boxplots of total sleep over 15 cycles (W = 860; *p* = 4.18e-07) and (G) of total number of long bout sleep episodes over 15 cycles (W = 264.5; *p* = 0.0005) show significant differences in the direction predicted by the model. Statistical comparisons were made using the one-tailed Wilcoxon’s test. * < 0.05, ** < 0.01, *** < 0.001, **** < 0.0001, and N.S. (Not Significant).

## Discussion

Through this manuscript, we show that an examination of the circadian regulation of sleep must involve the use of time-series analyses on sleep data rather than relying on locomotor activity data (Fig. 1). We have created tools to make such analyses accessible to a wider community of sleep researchers through an open-access R package [40] (different from our recently published MATLAB software, PHASE [73]; https://cran.r-project.org/web/packages/phase/index.html). By utilizing these tools, we derive a two-process model of sleep regulation for flies that is capable of accounting for the bi-episodic nature of fly sleep and the percentage of time spent sleeping under LD12:12 (Fig. 2). In addition, we show that ultradian rhythms in sleep are detectable in flies whose circadian sleep thresholds do not oscillate (Fig. 3). We then show that a two-process model in which the circadian clock speed and rates of the S-process do not covary is incapable of explaining *Drosophila* sleep rhythm strength and amounts across a wide range of periodicities under constant darkness (Fig. 4). However, covarying sleep homeostatic parameters along with the free-running period produces much better fits with experimental observations (Fig. 5) and suggests that circadian clock speeds are strongly correlated with the sleep pressure decay rate (Fig. 5). Furthermore, we show that our model predictions regarding the relationship between circadian period and amounts of sleep are in agreement with longer, deeper sleep bouts in flies, suggesting that short bouts are dispensable for capturing this relationship (Fig. 6). Using such longer bouts of sleep, we show that indeed *per^S^* flies exhibit several major properties of a homeostatic system that is characterized by faster decay of sleep pressure compared to wildtype flies (Fig. 7). These results underscore the utility of a formal two-process model for understanding the regulation of sleep in *Drosophila*.

### On ultradian rhythms in sleep and the inter-relationship between circadian period and rates of homeostatic processes

The theoretical framework around Borbély and Daan’s two-process models has played a critical role in the advances made in our understanding of mammalian sleep [2,4–7]. The conservation of sleep-like states across taxa and the power of *Drosophila* as a genetically tractable organism has made the fly a fruitful model system for the study of sleep regulation. However, the field of fly sleep has not benefitted from the strong theoretical framework provided by a formal two-process model. The appropriateness of this model as a framework for the investigation of fly sleep is exemplified by its prediction that ultradian sleep rhythms are a consequence of a functional S process in absence of oscillating thresholds (Figs. 3A and 3B). The two-process model of mammalian sleep has been adjusted to account for the presence of cycling NREM and REM sleep states in mammals, which produce ultradian EEG rhythms during bouts of sleep [74–76]. However, the ultradian changes in sleep state produced by normal sleep cycles occur during the circadian clock timed daily window of sleep and are therefore distinct from the ultradian rhythms in sleep bouts predicted by two-process models when the circadian cycling of sleep thresholds is abolished. Mistlberger and co-workers (1983) [77] detected the presence of ultradian rhythms in wheel running activity when the suprachiasmatic nuclei were bilaterally ablated, thereby abolishing circadian timekeeping. However, these ultradian rhythms were not explained in the context of the two-process model.

Previous work on loss-of-function *period* mutants detected the presence of ultradian rhythms in locomotor activity [47,48], which were hypothesized to be due to an uncoupling of ultradian oscillators [47,54,78]. However, our results, which demonstrate the presence of such rhythms in a range of loss-of-function clock mutants (Figs. 3D and 3E), suggest that they are a simple consequence of the rise and fall of sleep pressure between non-oscillating sleep thresholds, akin to an hourglass system as has been previously suggested [79] or a relaxation oscillator. It is important to note here that although the model predicts an ultradian sleep rhythm of ∼10-h, our results in clock mutant flies reveal much ultradian shorter periods, albeit with a wide distribution of periods that lack the stability of a robust circadian oscillator. Among the possible explanations for this discrepancy are a reduced distance between the thresholds in the mutants, a steeper rise or decay of sleep pressure, or non-oscillating but unstable thresholds in loss of function clock mutants. The large variance in ultradian periodicities that are detected may also be a consequence of independent stochastic fluctuations of the upper and lower thresholds and the distance between them. Indeed, a combination of all such factors may be at play. Moreover, enhanced sensitivity of *per^01^* mutants to sleep deprivation has been reported previously, suggesting a pleiotropic effect of the loss of *period* on the sleep homeostat [32]. Similar pleiotropic effects may also govern the S-process of other loss-of-function clock mutants, contributing to this large variation in detected ultradian periodicities. However, the quantifiable presence of such ultradian oscillations in sleep (Fig. 3) and the possibility of detailed characterization of their properties (Fig. 7) under conditions of non-cycling upper and lower thresholds promises to be a useful new approach for the study of the mechanisms regulating process S that are highly likely to complement work employing sleep deprivation to examine the homeostatic regulation of sleep.

The detectable effects of faster decay of sleep pressure in *per^S^*flies under LL conditions (Fig. 7) is remarkable because PER protein is constitutively low under continuous light [80]. This implies that there may be a fundamental relationship between clock speed and the rates of change of the sleep homeostatic system. It is possible that the free-running circadian period present during development shapes homeostatic systems to coordinate the build-up and/or decay of sleep pressure with clock speed. It is also possible that *period* alleles have different effects on the homeostatic control of sleep via epigenetic mechanisms. Recent studies have reported differential epigenetic regulation associated with cell populations with divergent periods and distinct levels of clock gene expression [81,82]. Our results in flies may be broadly relevant to animal sleep regulation, as similar results have been reported in humans [64]. Specifically, the human PER3^5/5^ polymorphism is highly enriched in morning chronotypes, and such early types also have faster-running circadian clocks. Remarkably, PER3^5/5^ individuals experience higher sleep pressures and a faster build-up of non-REM activity during sleep deprivation compared to another human PER polymorphism, PER3^4/4^ [64,83].

Previous studies that simulated S-process parameters in rodent models found considerable variation in the time constants of build-up and decay rates of the homeostatic sleep drive across species and among strains of the same species [5,46,84]. We compared these values and the time constants derived from human EEG data and found that rodent models have faster build-up and decay rates of the S-process compared to humans. We then looked at free-running periods of a commonly used mouse strain and that of humans and found that mice have faster running circadian clocks than humans [85,86]. This is consistent with our conclusion that speed of the circadian clock likely affects the rates of S-process increases and decreases. The faster build-up of sleep pressure than its decay in flies may be a necessary constraint (given the choice of circadian clock parameters chosen in this study) for facilitating two bouts of sleep/cycle in flies, where one bout is considerably longer than the other.

### The utility of the two-process model of fly sleep for understanding previous findings and the generation of new testable hypotheses

The integration of this quantitative model with previous work examining the effects of specific clock neuron types on sleep and the daily timing of neural activity they display, offers a unique and powerful framework for understanding the neural basis of circadian sleep regulation in the fly. An examination of our model in the context of the growing body of knowledge regarding the relationships between circadian clock neurons and sleep timing leads to specific and empirically testable hypotheses that have not emerged from current models. Furthermore, the roles of circadian clock neurons in sleep control have not been investigated in the context of cycling sleep thresholds.

Under normal circumstances (i.e., the absence of physiological or social challenges), our model suggests that the onset of daytime sleep is timed by the event of the S-process hitting the rising phase of the upper threshold (Fig. 8A). The onset of nighttime sleep, on the other hand, is timed by the falling phase of the upper threshold (Fig. 8A). Both, the morning, and evening bouts of wakefulness appears to be timed by the rising phases of the lower threshold (Fig. 8A). The falling phase of the lower threshold appears to support the consolidation and maintenance of nighttime sleep (Fig. 8A). Work over the last decade has assigned sleep- or wake-promoting functions to specific classes of circadian clock neurons (reviewed in [14]). For example, the small ventral lateral neurons (s-LNvs) and the dorsal lateral neurons (LNds) are thought to be wake-promoting, whereas the dorsal neuron classes are thought to be sleep-promoting (reviewed in [14,16]). In these cases, the daily timing of neural activity displayed by these cell types, as measured by relative Ca^2+^ levels, fits these proposed roles in sleep control well (Fig. 8A).

**Figure 8:**
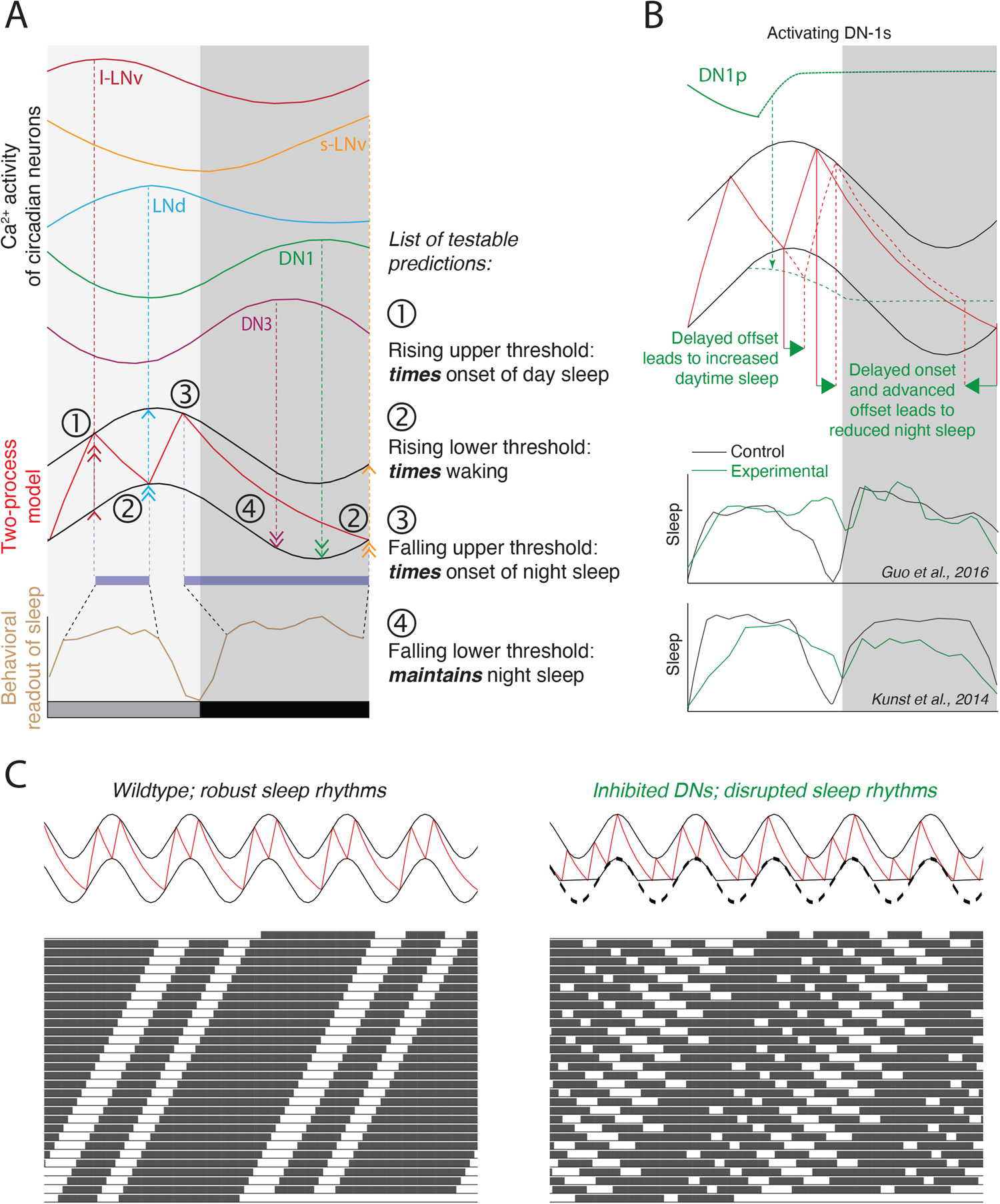
Utility of the two-process model in generating testable predictions. (A) Predicted roles played by the rising and falling phases of the thresholds in regulating the timing and maintenance of sleep, in the context of the two-process model. Hypothetical relationships between Ca^2+^ levels in circadian clock neurons, the two-process model, and sleep behavior. Data for the Ca^2+^ levels were extracted from Liang et al., 2016 [87] and smoothed using custom R scripts. Light gray shaded regions represent the subjective day phase, and the dark gray shaded regions represent the subjective night phase of one circadian cycle. The number of arrowheads is representative of the hypothesized strength of effects of calcium activity on the upper and lower thresholds, respectively. (B) A schematic representation of the effect of activation of DN-1s on sleep behavior previously reported by others. Exciting DN-1ps is predicted to produce an earlier fall and prolonged suppression of the lower threshold oscillations (dotted green lines). As a consequence of the earlier falling of the threshold, sleep pressure takes longer to hit the lower threshold (dotted red line) thereby increasing daytime sleep as was observed in two previous studies (bottom panel). The plots in the bottom panel are redrawn from Guo et al. (2016) [91] and Kunst et al. (2014) [89]. Green lines represent sleep in the DN-1p activated flies and the black lines represent unmanipulated controls. This manipulation also leads to an earlier termination of nighttime sleep as was reported by Kunst et al., (2014) [89], which is also explained by the effect of exciting DN-1ps on the lower threshold of the two-process model. (C) Two-process models and associated somnograms under unchanged conditions (left) and conditions of complete loss of decrease in the lower threshold (right). The complete loss of decrease in the lower threshold below baseline is taken to mimic a condition of inhibited DNs. Note the clear case of disrupted sleep/wake rhythms under constant darkness in this situation as compared to the wildtype situation (left). It is important to note that our specific quantitative predictions of sleep rhythms are produced by the best-fitting parameters of build-up and decay time constants derived from wildtype flies.

The s-LNvs display Ca^2+^ peaks that coincide with the morning bout of locomotor activity, the LNds display peaks that coincide with the evening bout of locomotor activity, and the DN-1 and DN-3s display their highest Ca^2+^ levels during the nighttime bout of sleep [87]. Integrating these patterns with our two-process model suggests that the s-LNvs control the timing of morning wakefulness by driving the initial rise of the lower sleep threshold, that the LNds promote the timing of evening wakefulness by the continued rise in the in the lower sleep threshold, and that the DNs promote the maintenance of nighttime sleep by driving the fall of the lower threshold throughout the night (Fig. 8A). Though this model explains daily sleep and wake patterns well, the timing and duration of daytime and nighttime sleep windows may also be controlled in other ways. For example, the distance between the two thresholds, the phase relationships between the two thresholds, or the relative amplitudes of the upper and lower threshold cycles might all be modulated by the circadian system to shape the timing and amount of sleep. These possibilities should be explored in future studies.

To test the utility of overlaying circadian clock neuron calcium activity and the two-process model for explaining previously published results and for generating new testable predictions, we considered the effects of DN neurons manipulation from previous studies. These studies have reached opposing conclusions regarding the role of the dorsal neurons 1 (DN-1) in sleep regulation: they have been proposed to be both wake-[88,89] and sleep-promoting [90,91]. Though these apparently incompatible conclusions may ultimately be explained by the different technical approaches used in these studies, the two-process model provides an explanation for how similar experimental manipulations of the same clock neuron classes might lead to contradictory conclusions. According to Liang et al., 2016 [87] the DN1ps display peak levels of neuronal activity in the middle of the night, a time associated with nighttime sleep, and a pattern that would appear to fit a sleep promoting role for these clock neurons. However, our model predicts that the effect of acutely exciting the DN-1ps would depend on the timing of the manipulation on neural signaling. For instance, if the DN-1ps promote the fall in the lower sleep thresholds, as suggested by our hypothesized relationship between the sleep thresholds and DN-1p Ca^2+^ profiles (Fig. 8A), exciting these neurons during the late afternoon would lead to an early and prolonged fall in the lower threshold (Fig. 8B). According to the model, this would increase daytime sleep by a delayed offset of daytime sleep and decrease nighttime sleep by causing a delayed onset and early offset of nighttime sleep (Fig. 8B). An increase in daytime sleep was used by Guo et al., 2016 [91] to support the conclusion that exciting DN-1ps promotes sleep. In contrast, Kunst et al., 2014 [89], present evidence for increased daytime sleep (Fig. 8B), but also describe a reduction in nighttime sleep, which, while not observed by Guo et al., 2016 [91], is predicted by our synthesis. Next, we asked what would happen to endogenous sleep rhythms if the DNs are inhibited. We model this by not allowing the lower threshold to fall below baseline (Fig. 8C). Our model predicts that inhibition of DNs would lead to disrupted sleep rhythms under constant conditions (Fig. 8C). Future work, employing time-series analysis of long, deep sleep bouts, will allow us to test numerous predictions provided by a synthesis of the two-process model and the daily patterns of clock neuron activity. This will provide a new approach to the examination of the circadian regulation of sleep in the brain.

### Speculations and Concluding Remarks

Our time-series analysis of *Drosophila* sleep detected the presence of ultradian periodicities under constant conditions in both wildtype flies and mutants lacking circadian timekeeping. Might these ultradian sleep rhythms reflect a cycling through different stages of sleep during the two daily bouts of sleep? Might such cycling occur at specific times of the circadian cycle? Answers to these questions await further detailed analyses of sleep time-series using the methods of analysis described here, all of which are freely available.

We have synthesized a formal, quantitative model of sleep regulation in *Drosophila* and have used it to guide the development of empirically testable hypotheses regarding the circadian control of sleep. We’ve also made this accessible to the wider sleep community by generating a Shiny-based GUI to facilitate simulations. We propose that this model can serve as a powerful framework for investigating the contributions of the various clock neuron to sleep regulation. The predictions made by this model were supported by experimental results, which were reminiscent of human studies on the relationships between circadian timekeeping and the homeostatic control of sleep. This suggests that the relationships between these two processes discovered in the fly will prove to be broadly relevant to our understanding of sleep regulation in animals. Our analysis also suggests that simple changes to the way we define sleep-like states will support new insights into the homeostatic regulation of fly sleep. Because the two-process model has been a major driving force for our understanding of sleep regulation in humans, we propose that adopting the two-process model to other genetically tractable experimental systems will support significant new insights into the molecular and neural mechanisms controlling sleep.

## Materials and Methods

### Fly husbandry

*Canton-S (CS*; BDSC stock number: 64349), *white^1118^* (*w^1118^*; [92]), *yellow, white* (*yw*; BDSC stock number: 1495), *period^TAU^* (*per^T^*; [65]), *period^SHORT^* (*per^S^*; [66]), *period^LONG^* (*per^L^*; [66]), and *period^01^* (*per^01^*; BDSC stock number: 80928), *timeless^01^* (*tim^01^*; BDSC stock number: 80930), *clock^JRK^* (*clk^JRK^*; BDSC stock number: 80927) and *cycle^01^* (*cyc^01^*; BDSC stock number: 80929) flies were used for the behavioral experiments reported in this manuscript. CS flies are also referred to as *period^+^* (*per^+^*) or wildtype interchangeably throughout the text, as all the *period* alleles used here were in a CS background. Flies were reared on Corn Syrup/Soy media made by Archon Scientific (Durham, North Carolina). Flies were approximately five days old on day one of all behavioral experiments.

### Generation of flies with a wide range of free-running periods

To generate flies with a wide range of period values, virgin females and males from all five *period* allele cultures (*per^T^*, *per^S^*, *per^+^*, *per^L^* and *per^01^*) were collected over the span of approximately five days. Reciprocal crosses were performed, resulting in a total of 25 distinct crosses. Male and female offspring from each of these 25 crosses, representing 50 distinct sex and genotype combinations (see Tables 1 and 2), were collected and assayed for locomotor activity as described below.

### Assay of locomotor activity

Locomotor activity was recorded using the *Drosophila* Activity Monitor (DAM) system (Trikinetics, Waltham, MA; https://trikinetics.com). Beam crossing counts were collected every minute for all experiments. Flies were recorded under LD12:12 cycles, constant darkness (DD) or constant bright white light (LL). In all experiments reported in this manuscript, flies were assayed under 25 °C and the intensity of white light was between ∼400 and ∼500 lux.

### Data analyses and visualization

#### Circadian time-series analysis of Drosophila sleep

Locomotor activity of approximately 32 each of CS, *w^1118^*and *yw* male flies was recorded under LD12:12 for nine days, after which activity was recorded under constant darkness (DD) for seven days. Beam crossing data from these recordings were used to determine the pattern and amount sleep for each fly. Individual fly locomotor activity and sleep time-series data, binned in 1-minute intervals, were subjected to (Chi-square) *χ^2^* or Lomb-Scargle periodogram analyses.

#### Calculating the percentage/amount of time spent sleeping

For fly sleep under LD cycles, the percentage of time spent sleeping was calculated by first computing the total sleep over the entire duration of the experiment for each fly (last four cycles of entrainment). This total was divided by the number of 24-h cycles (i.e., 4), and then multiplied by 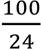
.

For flies under DD and for simulations of free-running sleep, the fly wise total amount of sleep (in hours) over the entire duration of the experiment or simulations was calculated. Because flies with different free-running periods would differ in the number of sleep/wake cycles experienced during free run – for example periods shorter than 24-h would complete more than 10 cycles in 10, 24-h days and clocks with periods longer than 24-h would complete fewer – we accounted for clock speed in our calculations. Therefore, all our estimates of sleep are hours of sleep/circadian cycle.

#### Continuous wavelet transforms as a tool to identify ultradian rhythms in sleep

Biological oscillations are often non-stationary time-series, i.e., their amplitude and period vary significantly over time. Such variations are often not of interest when estimating period and power of circadian rhythms. However, when attempting to identifying multiple periodic components from non-stationary time-series, traditional methods like the *χ^2^* periodogram often fail to accurately reflect the presence of multiple periodicities, especially when they are restricted to certain windows of time throughout the length of the time-series [41,49,51]. Recently, analytical tools to resolve circadian rhythms in both the time and frequency domain (i.e., provides period and amplitude information locally over a long time-series) have been developed and utilized [49–52]. We made use of one such tool, the continuous wavelet transforms (CWT), to visually examine periodic components in sleep time-series of our flies. We used functions from the WaveletComp package in R, which use a Morlet mother wavelet and resolves biological time-series in the time-frequency domain [49,93]. Complex-valued wavelet transforms such as the Morlet wavelet are useful because they provide amplitude and phase over time and generally preserve information about these locally [49,93]. To assess the presence of ultradian rhythms, sleep timeseries for all our genotypes for varying definitions of sleep were calculated in bins of 60-minutes. CWT were carried out and significance of detected periodicities were tested against a null hypothesis of “no periodicity” using shuffled timeseries. The null distributions were generated based on 1000 simulations of such shuffled timeseries. In order to quantitatively compare features of periodicities between genotypes (as shown in Figs. 3 and 7), we used a modified pipeline based on recent analytical developments [72]. We first performed discrete wavelet transforms (DWT) on 1-min binned sleep timeseries on our genotypes using a Daubechies wavelet of filter length 12 using the functions from the package “wavelets” on R [94]. We then computed the percentage of spectral variance accounted for by each level of the scale index as has been done previously [53,72]. Based on this, we identified three period bands of relevance in Fig. 3E (∼0.5-∼1-h, ∼1-∼2-h and ∼17-∼34-h) and Figs. 7C and 7D (∼1.07-∼2.1-h, ∼2.1-∼4.27-h and ∼4.27-∼8.53-h). We used 1-min intervals to compute the DWT to get the maximum resolution in identifying period bands corresponding to each wavelet detail (see Results).

In order to compare period values among genotypes within specific period bands (as in Fig. 7E), we revisited the identified periodicities from the CWT. In the interest of computation time, we performed the above-described continuous wavelet transforms on flies of CS and *per^S^*flies using 15-min binned sleep timeseries, and the significance of detected periodicities were based on 100 simulations of shuffled timeseries. We then computed median period and power for each fly in each of the three identified period bands after accounting for edge effects [72]. These values were then used to make between genotype comparisons. Statistical significance between genotypes were tested using bootstrapped *t*-tests with 10000 bootstrap replicates and were carried out using functions from the package MKinfer on R [95].

#### Time-series analysis of various definitions of sleep

Traditionally, *Drosophila* sleep has been defined as any bout of inactivity (usually the absence of beam crossings in DAM monitors) that lasts five minutes or more [12,13,15]. Though this method has been shown to overestimate the amount of sleep [96], its easy implementation and consistency over decades of research have made it an integral approach to measuring sleep in the fly. In addition to the traditional definition of sleep, we systematically changed the inactivity duration criterion for sleep to examine how the temporal regulation of sleep was affected by biasing our analysis to longer bouts of inactivity or specific ranges of inactivity. For example, a ten-minute inactivity duration definition of sleep would mean that any bout of inactivity lasting ten minutes or more would be considered sleep and subjected to time-series analysis, whereas a five to 20-minute definition of sleep would mean only bouts of inactivity that are more than five minutes but fewer than 20 minutes would be considered as sleep and subject to the analysis. These time-series were then subjected to Chi-square or Lomb-Scargle periodogram analyses, as described above.

### Figure generation

All figures were generated using custom scripts in R. We used the “wesanderson” package [97] to generate color palettes used in the figures, except in case of the acto-/somnograms, time-series plots in Fig. 1, the CWT spectrograms, and the Suppl. Figs. In these three cases, default color palettes were used to generate the plots.

#### Creating the two-process model and generating predictions

A major strength of the original two-process model was that it used EEG data to derive the parameter values of the sleep homeostat [4,5]. Because no equivalent physiological correlate of sleep is, as yet widely employed in flies, we relied on best-fitting approaches to find model parameters that best explain the waveform and amount of sleep in *Drosophila*. The basic structure of the two-process model used here is taken from a two-process model proposed by Daan et al., in 1984 [5]. The upper and lower thresholds are modeled as simple sinusoidal oscillations:

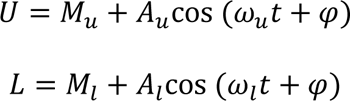

U and L are the upper and lower thresholds, respectively. *M*_*u*_ and *M*_*l*_ are the mean levels around which U and L oscillate. *A*_*u*_ and *A*_*l*_ are the amplitudes of the upper and lower threshold oscillations. ω_*u*_ and ω_*l*_ are the angular velocities of the oscillations. The angular velocities are determined by the period of the oscillation, which is expressed as ω = 2π/τ. τ is the period of the oscillation, which in this study, was fixed at the same value for both thresholds. φ is the phase angle by which the starting phase of the oscillations are adjusted and was likewise fixed at the same value for the upper and lower thresholds. The build-up of sleep pressure is modeled as:

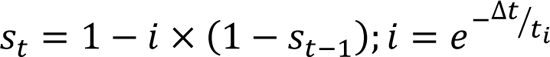

*S*_*t*_ is the value of sleep pressure at any given time *t*. *i* is the parameter determining the increase of build-up of sleep pressure. *S*_*t*−1_ is the value of sleep pressure at any given time *t* − 1. Δ*t* is the sampling interval (or bin size) for the simulations. This was set at 1-min because that is the resolution at which behavioral records were collected from our flies. *t*_*i*_ was the build-up time constant. The decay of sleep pressure is modeled as:

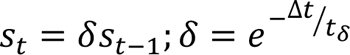

*S*_*t*_ is the value of sleep pressure at any given time *t*. δ is the decay rate of sleep pressure. *S*_*t*−1_ is the value of sleep pressure at any given time *t* − 1. Δ*t* is the sampling interval for the simulations.

*t*_δ_ was the decay time constant. The upper and lower thresholds were computed for the entire duration of the simulation. The simulations for sleep pressure build-up and decay were performed under the following conditions: at every time-step *t*, the program will increase sleep pressure as long as its value at the previous time-step (*t* − 1) was lower than the higher threshold at *t*. If at *t* − 1, the sleep pressure value is higher than the upper threshold at time *t*, the program will release the sleep pressure at *t*. Sleep pressure will continue to be released (i.e., fall) until it reaches the lower threshold value at *t*. The model was allowed to run for a duration of 60 days. The first 30 days of the simulation were discarded from analyses to remove potential transient behaviors. The remaining 30 days were used for all analyses. The window of time in the simulation where sleep pressure is being released is designated as the window of sleep. Using this window, the number of hours of sleep predicted by the model was computed for all conditions. This allowed for the estimation of percentage of time spent sleeping or the amount of sleep/circadian cycle. The sleep pressure time-series thus generated was subjected to *χ^2^* periodogram analyses to produced estimates of number of peaks in the periodogram and to assess the power of sleep rhythms. Power was defined as the height of the periodogram peak above the 0.05 significance cut-off. To facilitate easy access to predictions from these simulations, we have created a GUI which is available here: https://abhilashlakshman.shinyapps.io/twoprocessmodel/

#### Measuring sum of squared differences (SSD)

The S-process parameters that best recapitulated the waveform and amount of fly sleep were estimated using a sum of squared differences approach. For each combination of sleep pressure build-up (*t*_*i*_) and decay (*t*_δ_) parameters, we asked if the model produced two coherent peaks of sleep, and, if so, we assessed the difference between the % time spent sleeping predicted by the model and that of our experimental flies and this difference was squared to account for negative values. For the results reported in Fig. 2, SSD values were calculated across the three genotypes (using population averages; therefore, there are no estimates of errors on these parameter values) and, in the case of fitting reported in Fig. 5, SSD was computed for both the amount of time spent sleeping and for the normalized power of the sleep time-series periodogram. Given this approach, there is no good way to statistically distinguish neighboring points in the parameter space, except to choose the one with the minimum SSD. Any prediction that is close to empirical value would have small differences with the experimental data and consequently would have low SSD values. Therefore, the build-up and decay rates that together yielded the minimum SSD were considered the best-fitting parameters for our two-process models. See Suppl. Figs. 1, 3 and 6 for a guide to the algorithms for our model outputs. All our simulations were run on a 32-core AMD RYZEN Threadripper processor.

## Acknowledgements

This work was supported by a grant from the National Institute of Neurological Disorders and Stroke (R01NS077933) and start-up funds provided by the State of New York. We thank Michael Rosbash, Paul Hardin, Amita Sehgal, Patrick Emery, and Ralf Stanewsky for fly lines. We would also like to thank Aliya Fisher and Matthew Ciolkowski for technical support. We are grateful to Bill Joiner, Joydeep De, Meilin Wu, and Paul Franken for useful discussions of the work presented in this study. Finally, we thank Maria de la Paz Fernández, Matthew Ciolkowski, Budha Chowdhury, and Robert Veline for providing useful feedback on the manuscript. We would also like to extend our thanks to two anonymous reviewers for carefully reading a previous version of this study and providing valuable feedback.

