## Supplemental Tables, Figures, and Legends for "A two-process model of *Drosophila* sleep reveals an inter-dependence between circadian clock speed and the rate of sleep pressure decay"

*Running title: Drosophila two-process model*

**Suppl. Table 1:** Adjusted power values (mean±SEM) of *Canton-S* (*CS*), $w^{1118}$ and *yw* flies from Lomb-Scargle periodogram analyses on sleep and locomotor activity time-series under LD and DD conditions in the circadian range (between 16- to 32-h). Also reported are the differences in power values between sleep and locomotor activity powers for each genotype. Sample sizes are provided in parentheses.

| | ***CS*** | | $w^{1118}$ | | ***yw*** | |
|---|---|---|---|---|---|---|
| | **LD12:12** | **DD** | **LD12:12** | **DD** | **LD12:12** | **DD** |
| **Sleep** | 0.04 ± 0.01 (26) | 0.18 ± 0.02 (31) | 0.02 ± 0.00 (21) | 0.27 ± 0.02 (31) | 0.02 ± 0.01 (17) | 0.18 ± 0.02 (28) |
| **Locomotor activity** | 0.02 ± 0.00 (22) | 0.11 ± 0.01 (31) | 0.01 ± 0.00 (21) | 0.22 ± 0.02 (31) | 0.02 ± 0.01 (11) | 0.11 ± 0.01 (29) |
| **Δ Power** | 0.02 | 0.07 | 0.01 | 0.05 | 0.01 | 0.07 |

Step 1: Fix the amplitude of circadian oscillation of the thresholds and distance between them based on values reported in Daan et. al., 1984 which show bi-phasic sleep.

Step 2: Fix the $\tau$ of the circadian clock at 24-h.

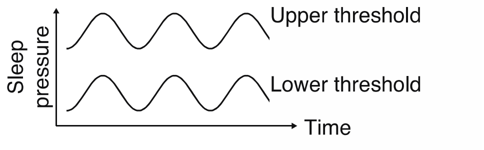


Loop 1: Vary time constant of build-up of sleep pressure (from 0.01 to 15 by 0.25).

Loop 2: Vary time constant of release/decay of sleep pressure (from 0.01 to 15 by 0.25).

Step 3: For each combination of build-up and release of sleep pressure create the homeostatic component of the model.

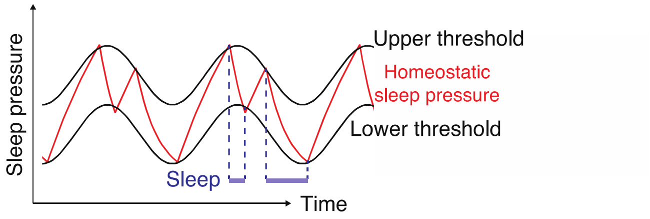


Step 4: Perform periodogram analyses and calculate % time spent sleeping for each such model.

Step 5: Compare these values with behavioral data to get an estimate of fit (SSD).

end loop

end loop

Step 6: Find out the combination of time constants of sleep pressure build-up and release time constants have minimum SSD.

**Suppl. Figure 1:** Sketch of the algorithm used for generating a two-process model for fly sleep.

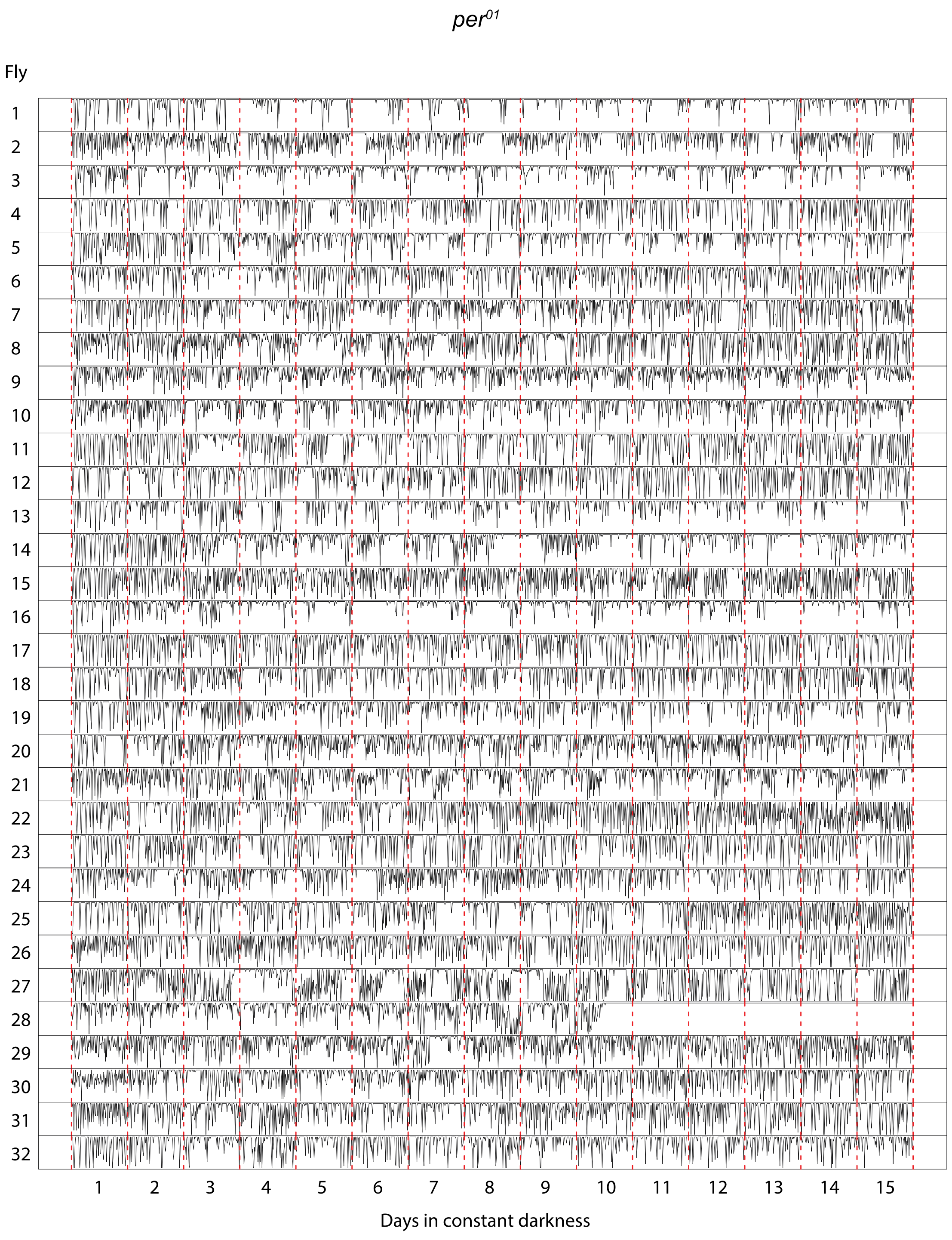


**Suppl. Figure 2:** Individual timeseries of *per*$^{01}$ flies under constant darkness. The red dotted lines demarcate days. Note the pervasive presence of ultradian rhythms in sleep with variable periodicities.

Step 1: Fix the amplitude of circadian oscillation of the thresholds and distance between them based on values reported in Daan et. al., 1984 which show bi-phasic sleep.

Step 2: Fix the rates of build-up and release of sleep pressure as identified from Fig. 2.

→ Loop 1: Vary free-running period of the clock (from 15 to 32-h).

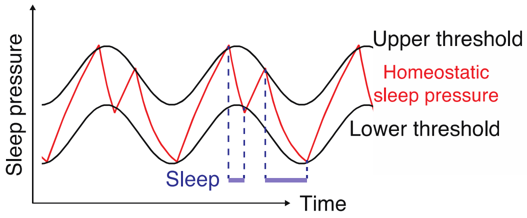


Step 3: In case of the model generated for each period value perform periodogram analyses and calculate amount of time spent sleeping.

end loop

Step 4: Generate plots to compare predictions from Step 3 and behavioral data for sleep.

**Suppl. Figure 3:** Sketch of the algorithm used for generating predictions from a two-process model with different free-running periods of the C-process.

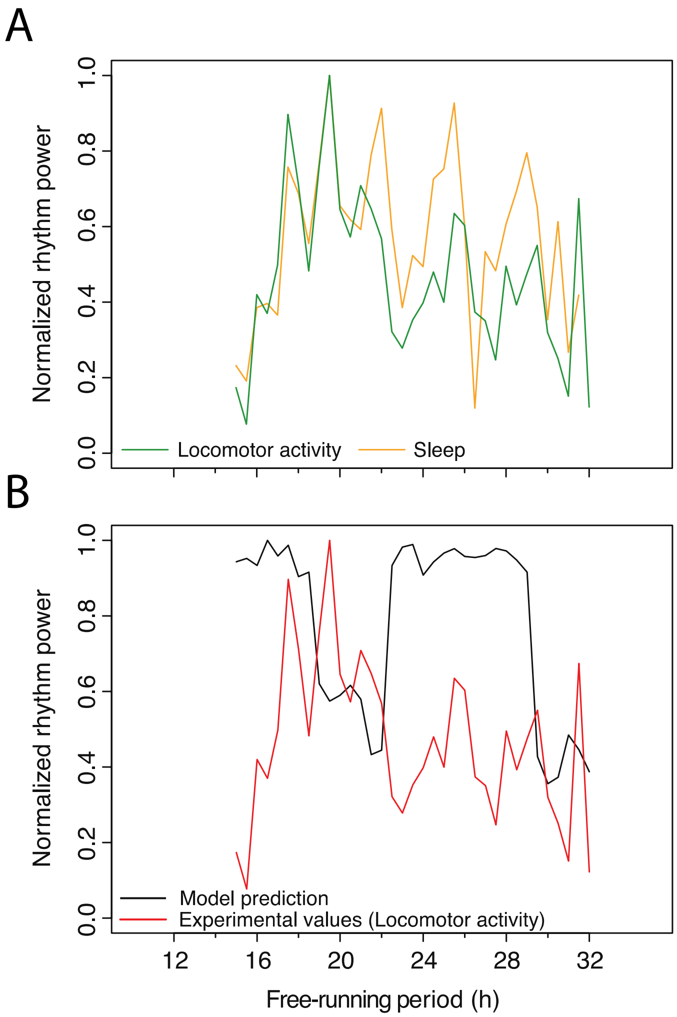


**Suppl. Figure 4:** (A) Median power of locomotor activity and sleep rhythms for flies with different free-running periods. (B) Median power of beam crossing rhythms across the entire range of free-running periods overlayed with the sleep rhythm predictions of the two-process model when only the free-running period was altered in the model.

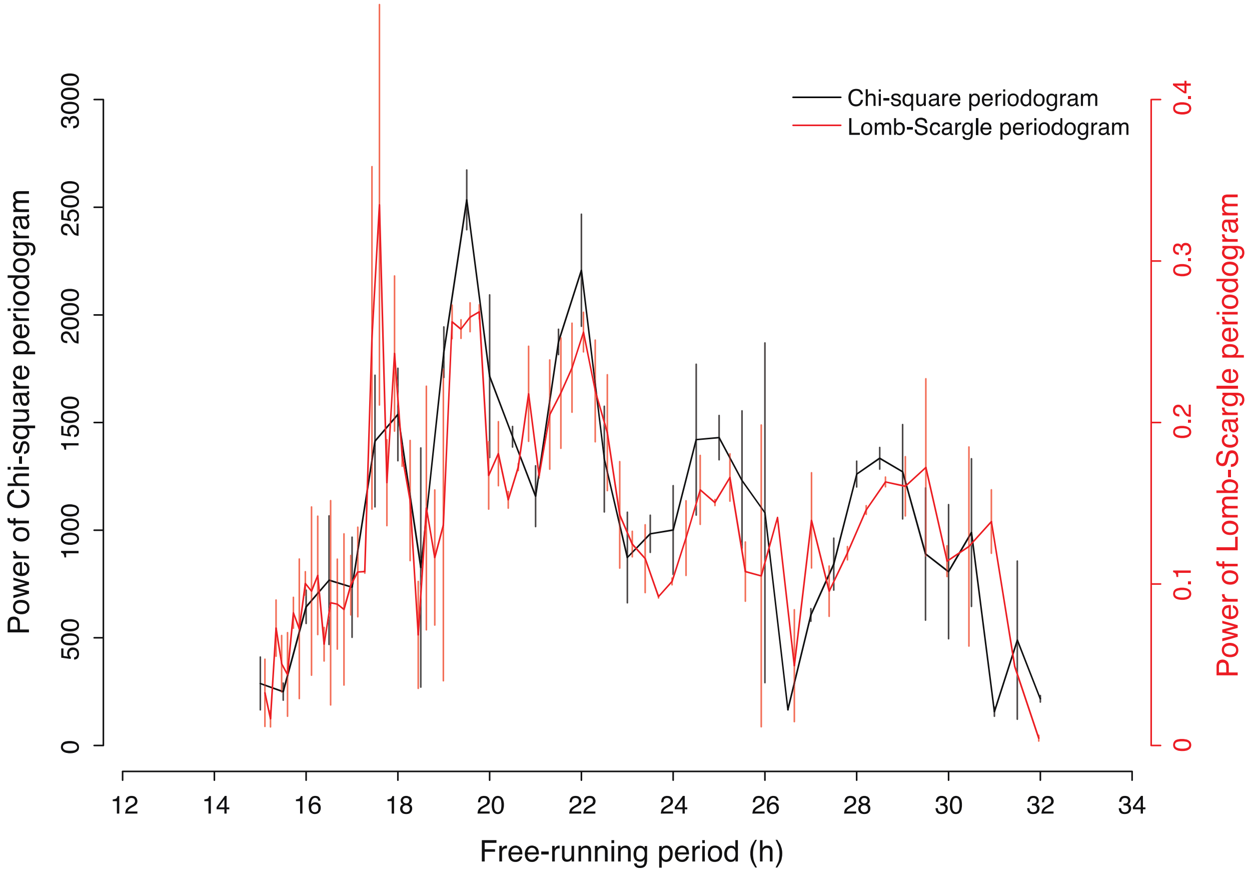


**Suppl. Figure 5:** Median power of sleep rhythms across the entire range of free-running periods using the Chi-square and Lomb-Scargle periodograms to highlight the fact that the relationship between free-running period and power of sleep rhythms is largely independent of the method chosen to estimate power of sleep rhythms. Error bars are SEM across two replicate experimental runs.

Step 1: Fix the amplitude of circadian oscillation of the thresholds and distance between them based on values reported in Daan et. al., 1984 which show bi-phasic sleep.

Loop 1: Vary τ of the circadian clock (from 15 to 32-h)

Step 2: Generate threshold oscillations for each free-running period.

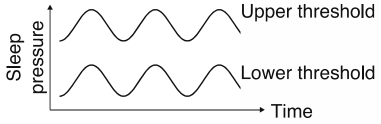


Loop 2: Vary time constant of build-up of sleep pressure (from 0.01 to 15 by 0.25).

Loop 3: Vary time constant of release/decay of sleep pressure (from 0.01 to 15 by 0.25).

Step 3: For each combination of build-up and release of sleep pressure create the homeostatic component of the model.

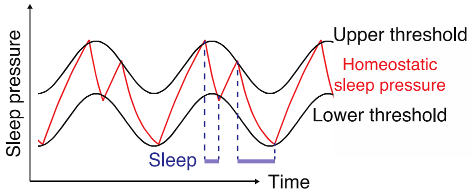


Step 4: Perform periodogram analyses and calculate amount of time spent sleeping for each such model.

Step 5: Compare these values with behavioral data to get an estimate of fit (SSD).

Step 6: Find out which combination of sleep pressure build-up and release have minimum SSD.

end loop

end loop

Step 7: Store the best-fitting sleep pressure parameters for each period value.

end loop

Step 8: Generate plots to compare predictions from Step 7 and behavioral data for sleep across different free-running periods.

**Suppl. Figure 6:** Sketch of the algorithm used for generating predictions from a two-process model with different free-running periods of the C-process when the S-process parameters were also allowed to covary.
